# Dopamine tracks adaptive learning of action representations

**DOI:** 10.1101/2024.07.28.605479

**Authors:** Maxime Come, Arnaud Lespart, Aylin Gulmez, Loussineh Keshishian, Joachim Jehl, Elise Bousseyrol, Steve Didienne, Eleonore Vicq, Tinaïg Le Borgne, Alexandre Mourot, Philippe Faure

## Abstract

Flexible decision-making requires not only updating values, but redefining which features constitute an action in a given context. We recorded nucleus accumbens (NAc) dopamine release while mice navigated a three-target intracranial self-stimulation foraging task in which outcomes were evaluated under three distinct reward delivery rules. Despite a constant motor repertoire, dopamine transients reorganized across contingencies and generalized linear models revealed context-dependent dopamine signal reflecting action direction, recent outcome-history, or target identity. Reinforcement-learning model comparison showed that these signatures are best explained by distinct reward prediction errors (RPEs) defined over different state–action representations, rather than a single fixed model-free scheme. A single deep reinforcement-learning agent trained by temporal-difference learning, recapitulated both the rule-specific policies and the corresponding dopamine signature. These results identify NAc dopamine as a dynamic readout of representation learning, remapping prediction errors onto the task features that define successful action as contingencies change.

## Introduction

Solving a puzzle often requires more than simply learning from outcomes—it also demands selecting which features matter depending on the context: A toddler might learn to sort objects by color in one game and by shape in another. This ability to flexibly reconfigure which environmental features are used to guide learning is fundamental to intelligent behavior. In computational terms, it requires not only the ability to learn action values from outcomes (reinforcement learning) but also to identify the relevant state space, i.e., the internal representation of the environment from which these values are derived. This process, known as representation learning, is increasingly recognized as a central challenge for neuroscience ^1–4^. While reinforcement learning theories formalize how agents update value expectations through reward prediction errors (RPEs), such mechanisms only operate effectively when the agent has selected an appropriate set of task-relevant variables. Critically, selecting the appropriate state representation allows animals to overcome a fundamental computational bottleneck known as the *curse of dimensionality* ^5,6^. In natural settings, choices often involve multiple features (e.g., spatial location, action history, outcome probability) that vary across contexts and time. Learning values for all feature combinations becomes intractable for standard reinforcement learning mechanisms, both because of the combinatorial explosion in possible states and because the mapping between features and reward can itself change ^7^. Flexible behavior therefore requires a mechanism to extract and update low-dimensional, task-relevant representations—effectively, learning *what counts as an action* in a given context—so that values can be learned efficiently.

Understanding how animals form and update such internal representations is essential for describing flexible behavior, particularly in environments with changing or ambiguous task structures ^3^. This issue has gained increasing importance as researchers shift their focus from experiments with a simple task structure to more elaborate tasks ^8–12^ that more closely resemble natural decision-making, with multiple (and possibly overlapping or competing) features that animals may use as state representations, as well as potentially abrupt changes over time in the state representations being used. Capturing the dynamics of representation change requires dedicated experimental and analytical approaches. While theoretical and behavioral studies have highlighted strategies to cope with high-dimensional environments^7^, identifying neural signals that reflect the internal representations used to guide learning and decision-making remains a major challenge. This identification can provide critical insight into which features are effectively used by animals to build a relevant internal model of the world — to predict values, compute errors, and guide goal-directed actions.

Dopamine (DA) is a very well-established substrate to signal value and compute reward prediction error (RPE) ^13–24^, integrating outcome-related dimensions in a common currency ^25–27^, and driving reinforcement learning and decision making ^19,25,26,28–31^. Because RPEs are necessarily computed within a specific state and action space, variation of DA transients may reveal which representation the animal is currently using to compute value. Recent work shows that DA can reflect inference over latent task states and context dependent expectation ^8,32^, as well as probability distributions over hidden causes ^33^, suggesting sensitivity to abstract structure beyond raw sensory features. However, direct evidence that DA dynamically reconfigures to track qualitatively different representations as animals shift between fundamentally distinct task contingencies remains lacking.

Here, we address this gap using a spatial decision-making task in freely moving mice. We imposed three reward rules ^9,10,34,35^ *i.e.,* deterministic, sequence-complexity and probabilistic, while keeping the same physical actions and outcomes, isolating changes in the computational structure of the problem from changes in motor demands. We combined this behavioral assay with fiber photometry recordings of DA release in the nucleus accumbens (NAc) and a two-level computational modeling approach. First, we used interpretable reinforcement learning models with hand-specified state and action features to formally compare alternative candidate representations and identify which features best explain NAc DA dynamics in each rule. Second, we trained a deep reinforcement learning (deep RL) agent with a simple feedforward architecture that learns its own internal representations from minimal inputs, and asked whether a single network can reproduce both the rule-specific behavioral policies and the rule-dependent modulation of RPE-like signals. This approach tests whether representation learning—extracting task-relevant features through error-driven plasticity—provides a parsimonious mechanism for the adaptive structure observed in DA signals. Together, these analyses show that DA signals dynamically reflect RPEs computed from context-specific features, revealing the internal model used by the animal in each condition, and that a deep RL agent trained by temporal-difference learning naturally recapitulates these rule-specific patterns under a fixed exploration policy. These findings position NAc DA as a real-time indicator of representation learning, providing a window into how the brain adaptively constructs and updates action-relevant structure in complex environments.

### Each reward context is associated with a specific reward-seeking strategy

Using different versions of a spatial bandit task adapted for mice ^9,10,34,35^, we aimed to obtain rule-specific and feature-dependent strategies. In this task, animals learned to navigate between three marked locations in an open field, each associated with an intracranial self-stimulation (ICSS) of the medial forebrain bundle (MFB). Mice could not receive two consecutive ICSS at the same location; and therefore, had to alternate between rewarding locations, resulting in a sequence of movements and binary choices (i.e., trials) **(Fig. 1A)**. Despite the apparent simplicity of this self-generated, goal-oriented behavior, mice can use different features of the environment to guide their actions and obtain rewards (**Fig. 1B**). Mice were initially trained in a deterministic context (*Det*) where all locations consistently delivered ICSS.

**Fig 1.**
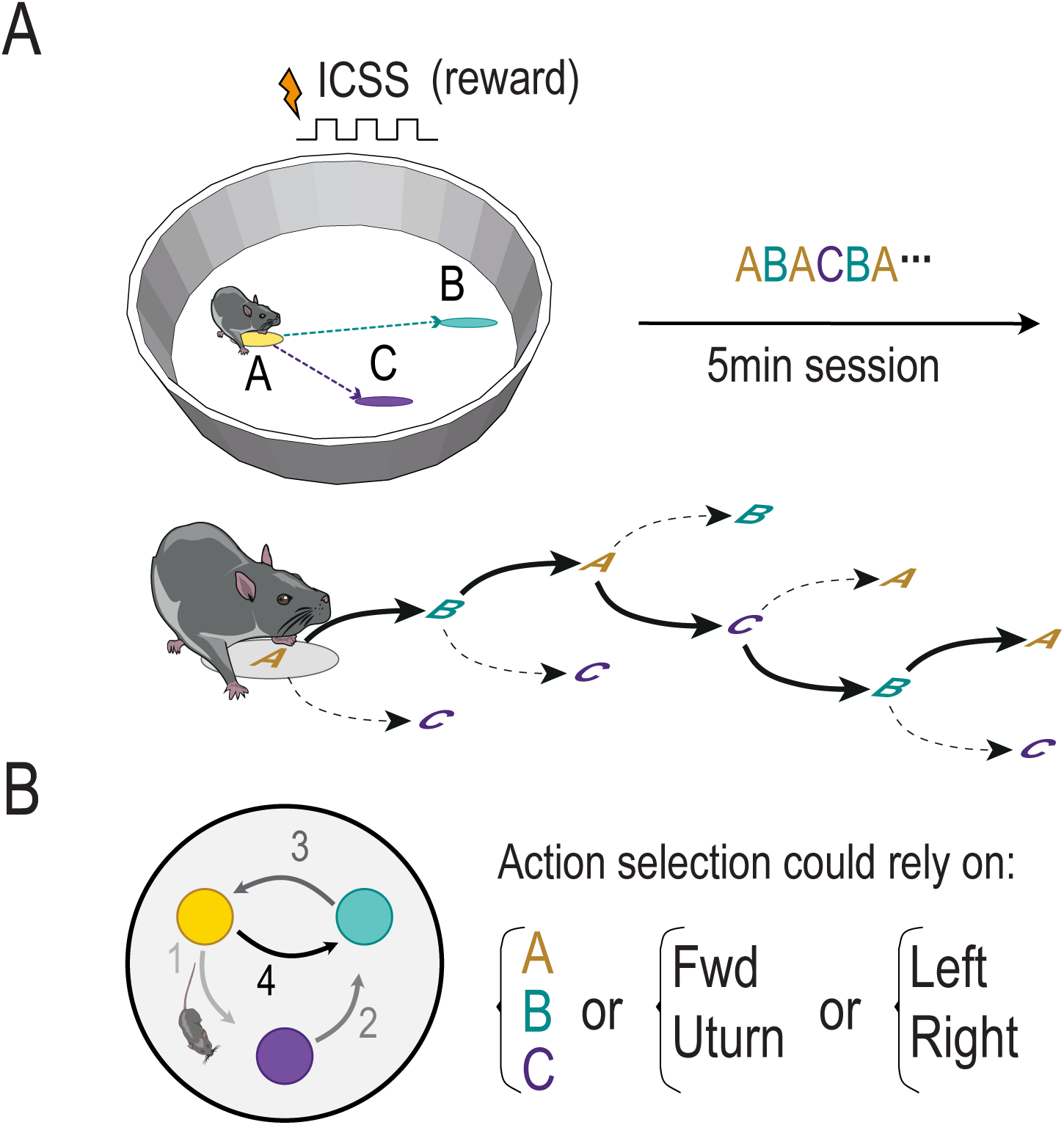
Decision making context: **(A)** Three explicit locations in an open field are associated with a possibility of intracranial self-stimulation (ICSS). Two stimulations could not be delivered consecutively in the same location, therefore animals learned to alternate between target. Mice perform successive binary choices to collect ICSS rewards. **(B)** Choice could rely on various overlapping sets of actions.

Although reward delivery was guaranteed at every location, this context nonetheless required animals to learn and internalize a simple structural rule: no two consecutive rewards could be obtained at the same location. As a result, mice had to alternate between the three available targets, producing sequences of choices constrained by spatial foraging dynamics. Initially, animals explored the environment with irregular trajectories and variable speeds. With training, they progressively optimized their behavior, developing stereotyped circular trajectories with minimal U-turns and ballistic movement profiles **(Fig. 2A)**. These behavioral adaptations reflect the acquisition of a minimal internal model of the task structure, allowing animals to predict that remaining at the same location would not yield further reward. This is further supported by the progressive increase in the number of trials **(Fig. S1A)**, reflecting improved efficiency in task execution throughout learning. Thus, although the Det context does not require complex credit assignment or value updating, it provides an essential baseline for evaluating the subsequent emergence of more sophisticated decision strategies when the reward structure becomes ambiguous or non-deterministic **(Fig 2B, Fig S1B-C)**.

**Fig. 2.**
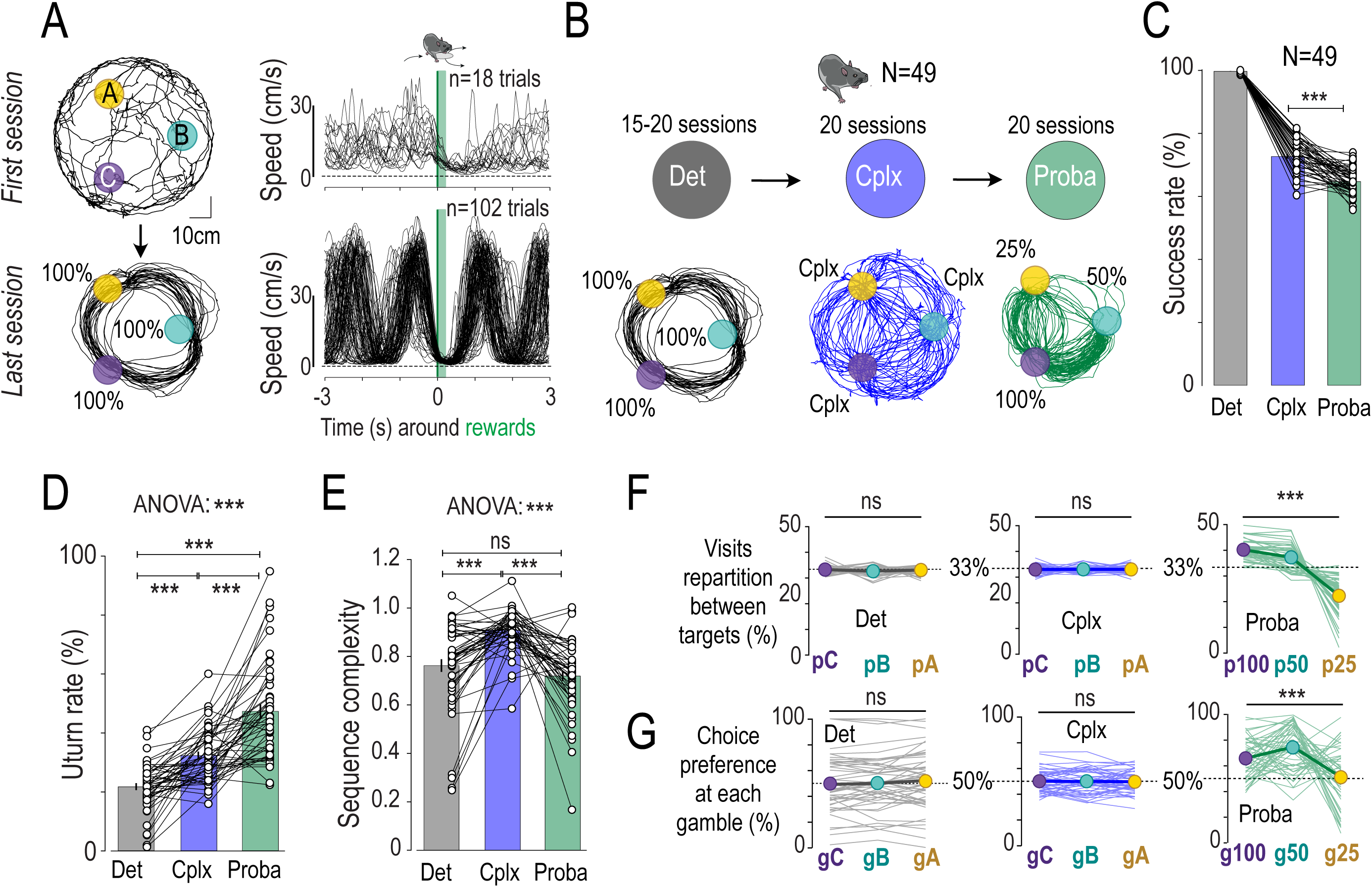
Mice display distinct reward seeking strategies adapted to each rule. **A.** (Left) Trajectories throughout conditioning and (Right) Speed profiles around ICSS in the deterministic context (First and Last sessions). **B.** Three reward delivery rules were successively proposed: Deterministic (*Det*) where all trials were rewarded (P=100%), Complexity (*Cplx*), where trials are rewarded based on sequence variability, and Probabilistic (*Proba*), with each target associated to a given probability (P=25%, 50%, and 100%). **C-E** Comparison of success rate, sequence complexity and Uturn rate reveals distinct reward seeking strategies across contexts. **F.** Locally, a mouse on one location (ex: p_A_) has the choice between the two other points (ex: p_B_ vs p_C_), and therefore performs a gamble computed as *g_A_ = P(p_B_|p_A_)*. g>50% corresponds to clockwise rotation for *Det* and *Cplx*, and to preference for highest probability of reward for *Proba*. Proportion of target visits and choice preference at each gamble show a bias for circular foraging in Det, exploitation in *Proba*, and randomness in *Cplx*. Data are shown as individual points, and mean ±sem. N=49 mice (23 males and 26 females).

In the complex context (Cplx) **(Fig 2B** middle), reward delivery was determined by the variability in recent choice patterns (see methods and **Fig S2A)**^10^, computed relative to the last nine decisions using a complexity algorithm. This rule requires animals to overcome a key limitation of model-free learning strategies. In contrast to the Det and Proba rules, where individual actions or spatial locations could be associated with consistent reward outcomes, the Cplx rule imposes no such direct contingency. Instead, animals must maximize variability in their behavior over time, independently of any fixed spatial preference or repeated motor strategy. This configuration renders previously learned policies a priori ineffective, and creates a setting in which trial identity loses its predictive power **(Fig S2B)**. Importantly, solving the task does not require tracking a large set of state-action pairs, which would be computationally costly, but rather adopting a *compressed representation* of the task, in which increasing sequence variability becomes the feature associated with reward ^10^. As such, the Cplx rule constitutes a strong test of representation learning, assessing whether animals can adapt their internal model when the structure of reward contingencies is non-trivial and abstract.

In the probabilistic context (Proba), each location was associated with a fixed probability of reward delivery (p100, p50, or p25) **(Fig. 2B** right). Unlike the Cplx rule, this setting involves stochastic outcomes and requires animals to integrate probabilistic feedback over time to optimize their choices. Successful performance relies on estimating expected values for each location and adapting spatial preferences accordingly. This context thus engages value-based decision-making based on feature-specific learning, with spatial location becoming the task-relevant dimension. In contrast to Cplx, where individual outcomes are largely uninformative without reference to prior behavior, the Proba condition allows a more classical reinforcement learning strategy to emerge — but still requires flexible updating of value representations as animals recover from the behavioral variability induced by the previous rule ^35^.

These three distinct reward structures resulted in distinct trajectory patterns **(Fig 2B)**, success rates **(Fig 2C)**, and decision-making strategies **(Fig 2D-G)**. In *Det*, animals tended to adopt circular trajectories with minimal U-turns (∼20%) **(Fig 2D)**. In contrast, the *Cplx* rule resulted in random trajectory patterns characterized by high sequence complexity (**Fig 2E, Fig S2B)**. In *Proba*, mice exhibited a bias toward locations with higher probability of reward delivery **(Fig 2G)**, resulting in a high percentage of U-turns (**Fig 2D**) and a preference for p100 and p50 (**Fig 2F**). We ensured that those differences in decision strategy were not due to motivation or vigor to perform the three versions of the task **(Fig S3A-B)**. Overall, while the basic design of the task remained constant, each rule instantiated a distinct causal relationship between actions and rewards. The evolution of behavior across contexts demonstrates that mice are capable of extracting and adapting to these contingencies, dynamically modifying their decision strategies. This makes the task well-suited to track both behavioral flexibility and its neural correlates over time, in particular through the lens of DA dynamics and internal representation learning.

### Nac Dopamine dynamics reveal expectations built upon rule-specific features

We next examined DA release dynamics during the task, across the three rules, using the fluorescent sensor GRAB_DA2m_ ^36^ expressed in the lateral shell of the nucleus accumbens (NAc) in a new cohort of wild-type male mice (**Fig 3A, Fig S4A**). Positive events in DA release occurred upon receiving expected rewards, whereas negative events were observed when expected rewards were omitted **(Fig 3B-C, Fig S4B)**, indicative of a negative RPE (for simplicity, these events, whether positive or negative, are referred to as transients). Similar responses were observed while recording Ventral Tegmental Area (VTA) DA neurons activity with GCaMP in DAT-iCre mice, ensuring consistency in the interpretation of DA dynamics between release and somatic processes **(Fig S4C-D)**. Analysis of the amplitude distribution of DA transients (positive and negative) across the different rules showed greater variability compared to unexpected random stimulation in a rest cage, suggesting an active mechanism related to reward expectation modulating the DA response, rather than being a mere response to the ICSS (**Fig 3D**). This modulation is particularly important given that the ICSS directly activates MFB fibers, raising the possibility of DA release being directly driven by stimulation. The observed variability across task contexts thus strongly supports a dynamic, expectation-dependent component in DA signaling that add to the electrical stimulation signal. Additional experiments with unexpected rewards delivered either during the task but off-target (i.e. when the animal was in-between rewarded locations, **Fig 3E**) or in a rest cage (**Fig S5A-D**) demonstrated a larger response compared to expected rewards during the task, yet only after conditioning, further supporting the role of expectation in modulating DA release. We also controlled for a potential impact of sensor fatigue and found no effect on DA signal when stimulations were given in the rest cage with varying durations in-between stimulation (matching those observed in the task, typically from 2s to 7s) **(Fig S5E-F)**. Altogether, these findings confirm that DA positive transients during the task are not merely driven by the delivery of MFB stimulation, but are modulated by reward expectation. The amplitude of DA signals varied depending on whether outcomes were expected or not, and this modulation differed across contexts and over time. These results are consistent with the notion that DA release reflects reward prediction error (RPE) signals, with positive transients following unexpected rewards and negative transients after unexpected omissions. Notably, similar ICSS pulses evoked different responses depending on behavioral context—highlighting that DA signals are influenced by ongoing task engagement and learning history, rather than being reflexive responses to stimulation alone. These observations support the use of DA transients as dynamic markers of outcome expectation. To better understand the basis of these expectations, we next asked: what task features are DA signals sensitive to, and how does this sensitivity vary across rules? To do this, we applied generalized linear models (GLMs) to analyze fluctuations in DA transients amplitudes across trials, running separate regression analyses for each individual mouse at the end of each rule (last two sessions) **(Fig 4A)**. The predictors included current and previous trial outcomes (P: reward or omission), the specific target where outcomes occurred (T: locations pA, pB, and pC; or p100, p50 and p25 in Proba), the direction taken (UT: Forward movement or U-turn) and side (RL: left or right choice). These variables were selected based on their relevance to the specific constraints of each task rule, and their potential to capture the different features that mice could use to build internal models of the environment. In the *Det* setting, where all trials were rewarded, we observed that the key predictor for differentiating trials was direction (UT) but not side nor target **(Fig 4B).** In the *Cplx* setting, trial outcome accounted for the biggest part of DA variation (positive for rewards, negative for omissions, **Fig 4C**), with an additional positive effect of previous outcome (having received an omission at trial n-1 increases DA signal at trial n), regardless of targets, side or directions. In *Proba*, this effect of previous outcome disappeared, and the target probability (T) significantly influenced DA variations **(Fig 4D)**.

**Fig. 3:**
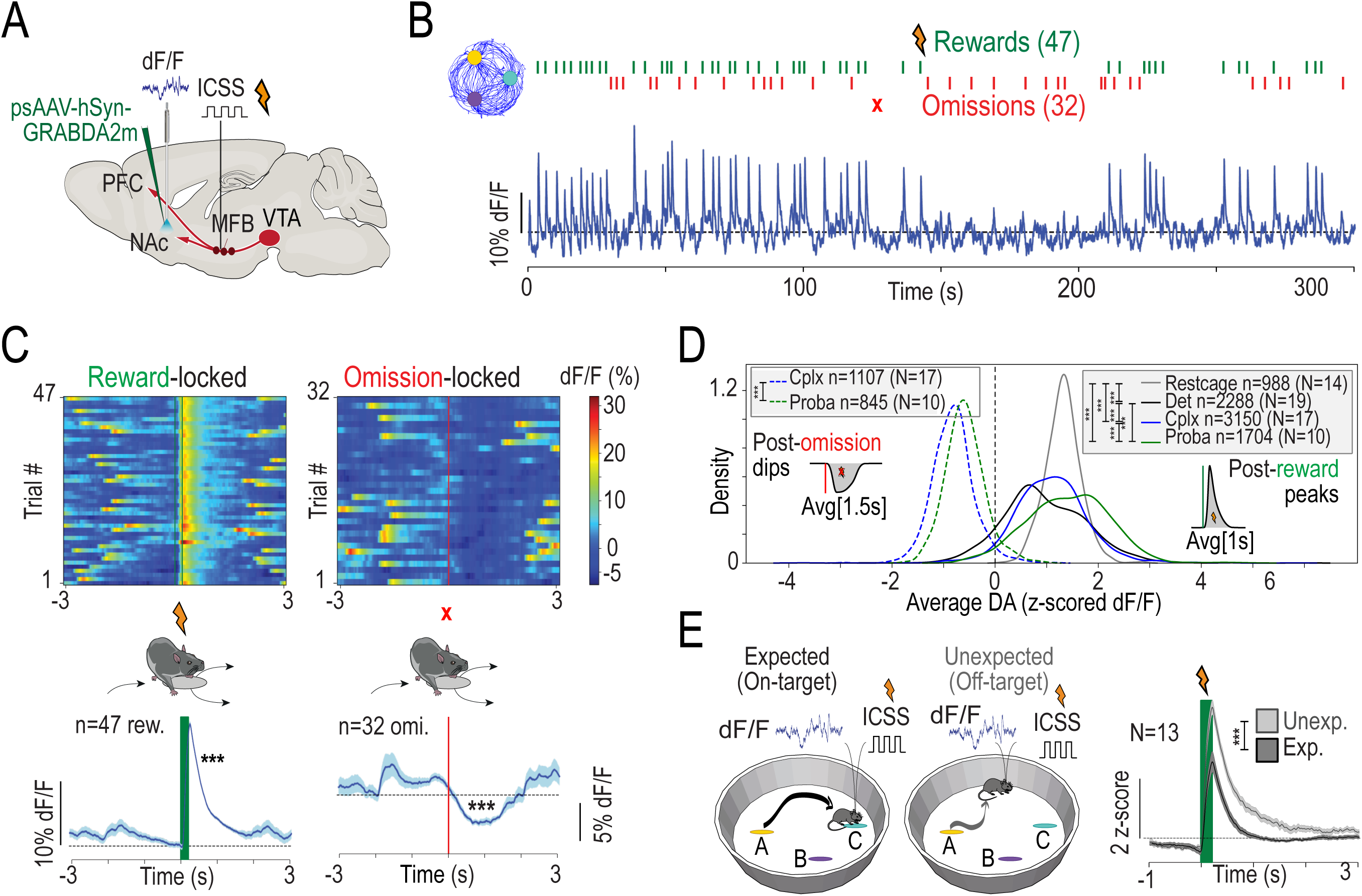
NAc DA release dynamics reveal expectations built upon rule-specific features. **A.** Schematic of the experimental design to record DA release during the task with chronic fiber photometry. **B.** Representative signal from one 5-min session with rewards (green) and omissions (red). **C.** For the same example session, signal is time-locked on location entry (t=0) and averaged. Rewards induce peaks and omissions induce dips of DA release. **D.** Density distribution of averaged DA variations (Peak) for rewards and omissions for the last two sessions of Det, Cplx or Proba, and for random stimulations in the rest cage (performed on last day of Det). **E.** After conditioning, mice were randomly and unexpectedly stimulated during the task outside of the rewarded zones (off-target), triggering DA peaks of greater amplitude.

**Fig. 4:**
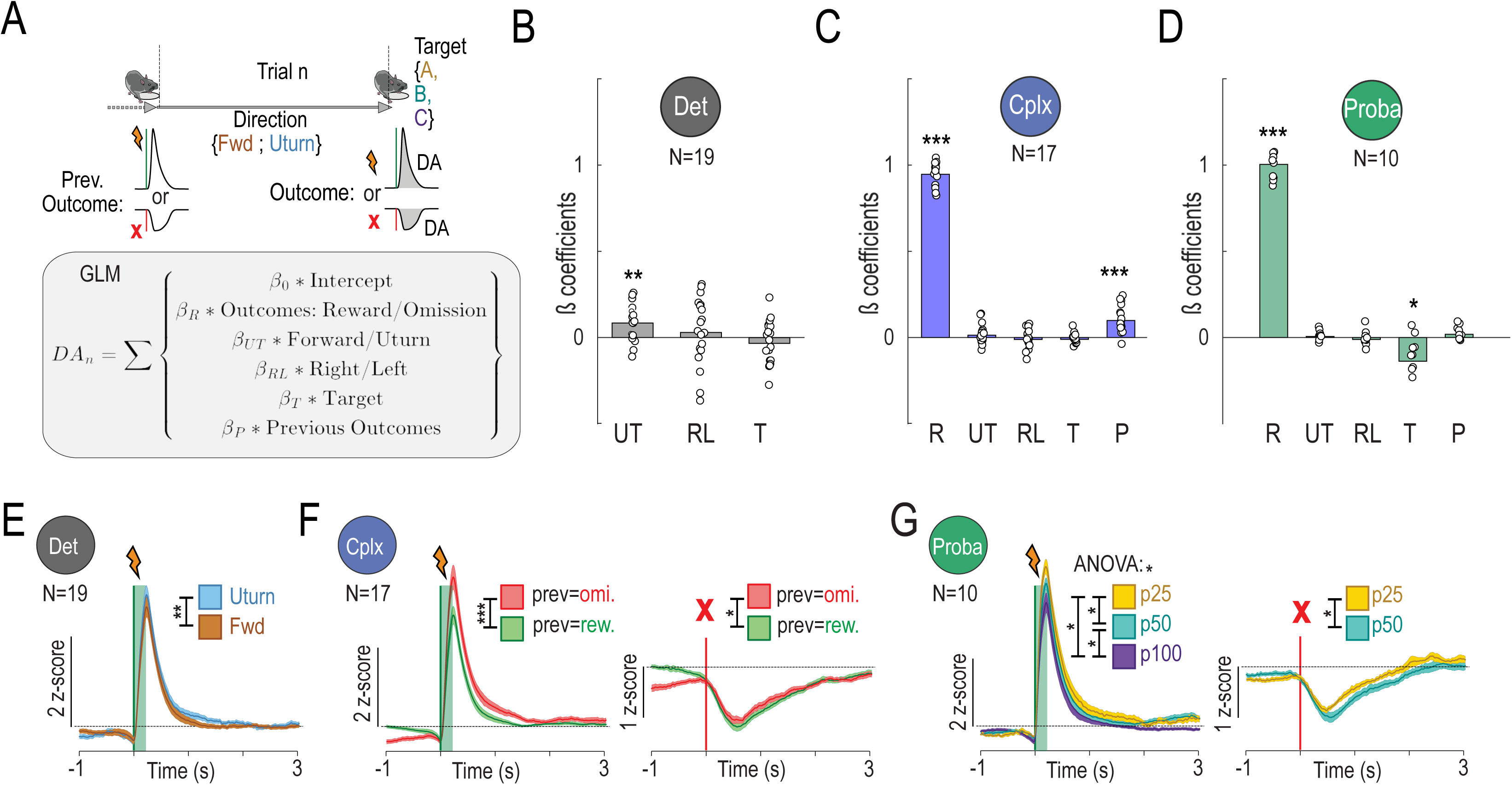
Defining sources of DA variations. **A** Each trial is defined by predictors (outcome received, previous outcome received, trajectory chosen to reach target, and target chosen) to fit DA amplitude using GLMs. **B-D.** GLM results at the end of Det, Cplx and Proba. Features explaining DA variations vary across contexts. **E-G.** Direct analysis of DA transients locked on those significant features (Uturn vs Fwd in Det; reward vs omission at previous trial in Cplx; p25 vs p50 vs p100 in Proba). Data are shown as individual points, and/or mean ±sem. n is the number of trials, N the number of mice in each condition.

Overall, the GLM analysis revealed that the primary drivers of DA fluctuations changed depending on the task setting, with direction, trial outcome, and target probability each playing distinct roles. Direct examinations of DA transients, categorized by direction, previous outcome or target, supported the strength of these effects. In *Det*, DA release varied with the direction **(Fig 4E, Fig S6A)** but not with the target (**Fig S6B**). In *Cplx*, omission on previous trial led to greater reward-induced and shallower omission-induced transients (**Fig 4F, Fig S6C**), while neither the target nor the direction showed significant effects (**Fig S6D-E**). At the end of the *Proba* setting, the DA signals were negatively influenced by target probability, with higher probabilities resulting in smaller positive transients for rewards and deeper negative transients for omissions **(Fig 4G, Fig S6F)**. Finally, no effect of direction was observed on DA transients **(Fig S6G)**, and regarding outcome at previous trial, we observed a small effect only for rewarded trials **(Fig S6H).** Altogether, these results reveal specific patterns in the modulation of DA transients across task settings. Notably, DA fluctuations were not tied to the same task features across rules: outcome history dominated in Cplx, direction in Det, and target identity in Proba. This dissociation indicates that the features driving DA transients are not fixed, but flexibly selected depending on task structure—supporting the view that phasic DA signals adapt to the relevant contingencies of each context.

### NAc dopamine cannot be explained by single model-free reinforcement learning but encodes state-specific RPEs

GLM analyses demonstrated that DA transients are modulated by distinct task features depending on the current rule—such as direction in Det, outcome history in Cplx, and target identity in Proba. However, they do not clarify *how* such features emerge nor whether the signal reflects a genuine reward prediction error (RPE). To directly test the RPE hypothesis, we used reinforcement learning (RL) models to generate trial-by-trial RPEs from the actual sequences of choices made by the animals (**Fig S7)**. A critical aspect of RPE computation is that it depends on the internal representation used to track expected value and on a well-defined set of competing actions with distinct consequences: from a given state, each choice must correspond to a different state transition (i.e., reaching a given location, such as pA, is uniquely determined by one action from that state, rather than being attainable via multiple equivalent actions). For instance, RPEs can be derived from predictions based on action direction (e.g., U-turn vs. forward), spatial target (e.g., pA vs. pB), or simply trial outcome regardless of action or location. These different assumptions correspond to different candidate representations of the task structure. By comparing theoretical RPEs from each model to recorded DA transients, we can infer which representation best matches the neural data in each context.

Based on our experimental results, we tested four models reflecting distinct representational schemes (**Fig. 5A**). Model 1 (M1) used a trial-based representation in which all choices updated a single global value, ignoring action or target identity. Model 2 (M2) assigned separate values to U-turn and forward actions, capturing a motor-based distinction. Model 3 (M3) tracked expected value for each target location independently. Finally, Model 6 (M6) implemented a non-abstracted, fully specified model-free scheme in which values were associated with each unique state-action pair (e.g., target × direction), reflecting learning without any structured generalization. All models were implemented using the Rescorla-Wagner rule to generate trial-by-trial RPEs, which were then used to assess their ability to reproduce key features of the DA signal in each task context.

**Fig. 5:**
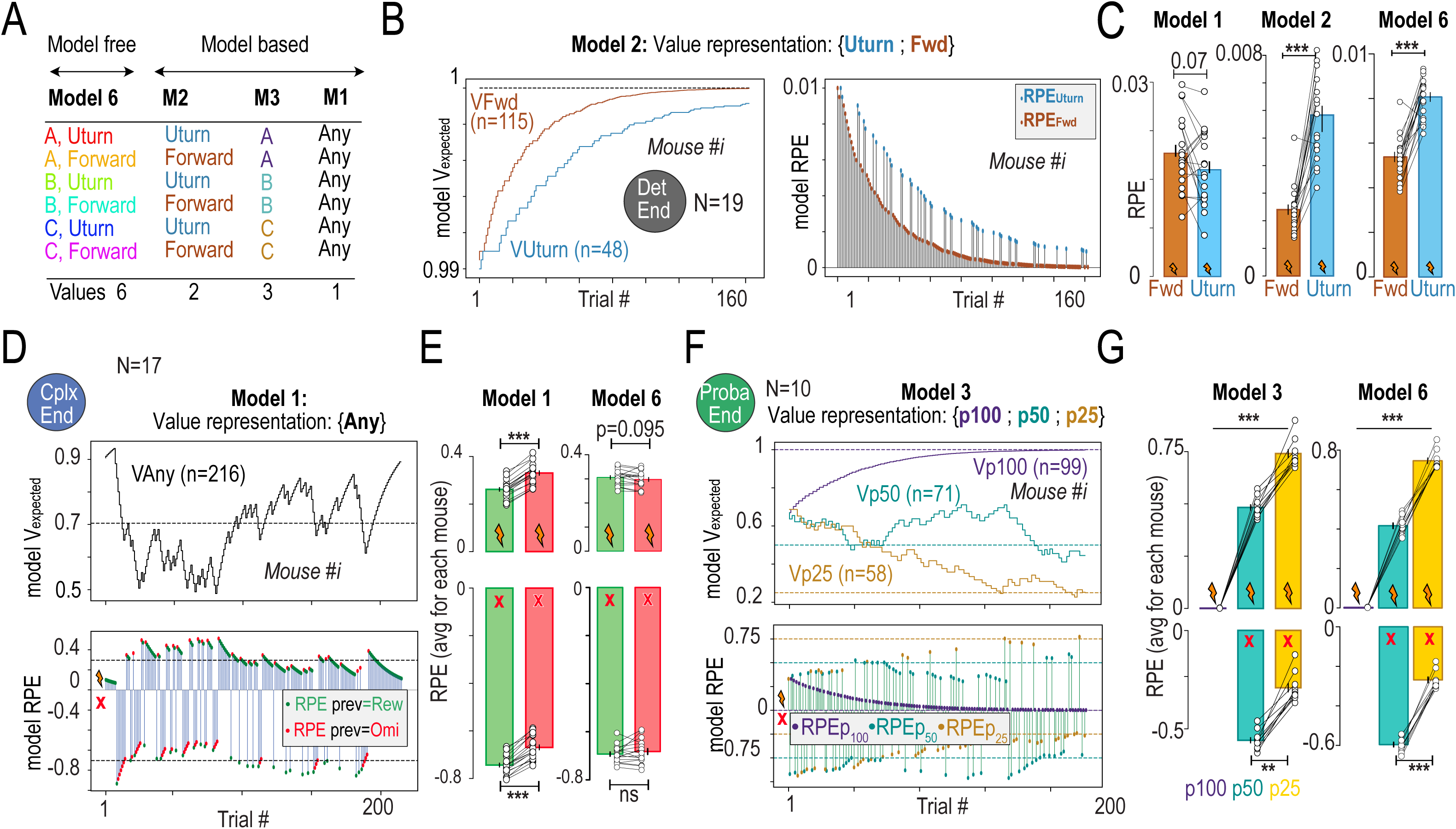
DA signal embeds an RPE component, modelled from distinct value representations specific to each rule. **A.** Mice choice sequences were taken to train Reinforcement Learning (RL) algorithms, testing three possible action representations to update values and compute corresponding RPEs. Model 1 (M1) treats all trials equally with fluctuating { V_Any_ }. M2 updates a set of two distinct values { V_Fwd_; V_Uturn_ }. A spatial model (M3) computes three independent values for each target { V_pA_; V_pB_; V_pC_ }. M6 computes 6 independent value considering forward and Uturn for each position A, B or C. We then trained another GLM assuming DA = V_obtained_ + RPE, with trial RPEs generated from M1, M2, M3 and M6. **B-G.** Models reproducing RPEs that explained DA variations vary across contexts. **B.** In M2-Det, convergence toward 1 is slower for Uturns, leading to higher RPE_Uturn_ and reproducing DA data. **C.** M2 and M6 reproduce RPE data but not M1**. D.** In M1-Cplx, V_Any_ is always updated and fluctuates around mean success rate. Plotting corresponding RPEs regarding current and previous outcomes mimic DA data in M1 but not M6. **F-G.** In M3-Proba, value of each target converges and then fluctuates around its probability, and corresponding RPEs reproduce DA data in M3 and M6. Data are shown as individual points, and/or mean ±sem. n is the number of trials, N the number of mice in each condition.

In the Deterministic (Det) context, mice developed a directional bias in their choices, favoring forward movements over U-turns, which was mirrored by asymmetries in DA transients (**Fig. 4E**). Consistent with this, models M2 and M6 produced direction-dependent RPEs matching the fiber photometry data (**Fig. 5B–C**), whereas M1 and M3 did not (**Fig. 5C, Fig. S8A**), with M1 even predicting the opposite asymmetry. This supports the idea that, in Det, DA fluctuations reflect value representations incorporating action direction. In the complexity (Cplx) rule, where rewards depended on variability in recent choice sequences, DA transients were modulated by recent outcome history with no dependence on action or spatial features (**Fig. 4F**). Only the trial-based model M1 captured this pattern (**Fig. 5D–E**), whereas models with directional or spatial structure introduced effects not supported by the data (**Fig. 5E, Fig. S8B**), suggesting that animals relied on a simplified, history-tracking representation under Cplx. In the probabilistic (Proba) context, mice progressively developed spatial preferences aligned with fixed reward probabilities, and DA transients showed corresponding target-specific modulation (**Fig. 4G**). Models M3 and M6 reproduced these graded RPE differences across locations (**Fig. 5F–G),** although both tended to introduce additional modulations; among these, only the outcome-history effect (and not direction) was supported experimentally (**Fig. S6H** vs **Fig. S8C**).

Together, these results show that the DA signal does not reflect a unitary, fixed model-free mechanism (M6), but rather encodes context-specific RPEs based on dynamically selected internal representations. This flexible alignment between task structure and neural computation supports a key role for dopamine in representing the features that matter most for decision-making in each context.

### A deep RL model captures behavior and Nac dopamine signals

To understand the computational principles that could jointly generate the behavioral trajectories and DA signals observed across rules, we next turned to a deep reinforcement learning (deep RL) approach. Rather than prescribing a specific representation a priori (trial-based, direction-based or spatial), we asked whether a simple neural network trained by RPEs could *learn* appropriate internal representations and reproduce both behavior and DA-like signals. We implemented a minimal feedforward network with one hidden layer trained by temporal-difference learning (Methods, **Fig. 6A**). On each trial, the network received the animal’s current location (A, B, or C) and produced five action values (Q-values): two motor actions (Left, Right) and three target-specific actions (go to A, B, or C). Because motor and spatial descriptions can be partially redundant (e.g., “Left” and “go to A” may lead to the same choice), outputs are not mutually exclusive. The model sampled actions from a softmax over Q-values, received reward or omission according to the current rule, and computed a scalar RPE from obtained versus predicted outcome for the chosen action (Methods).

**Fig. 6:**
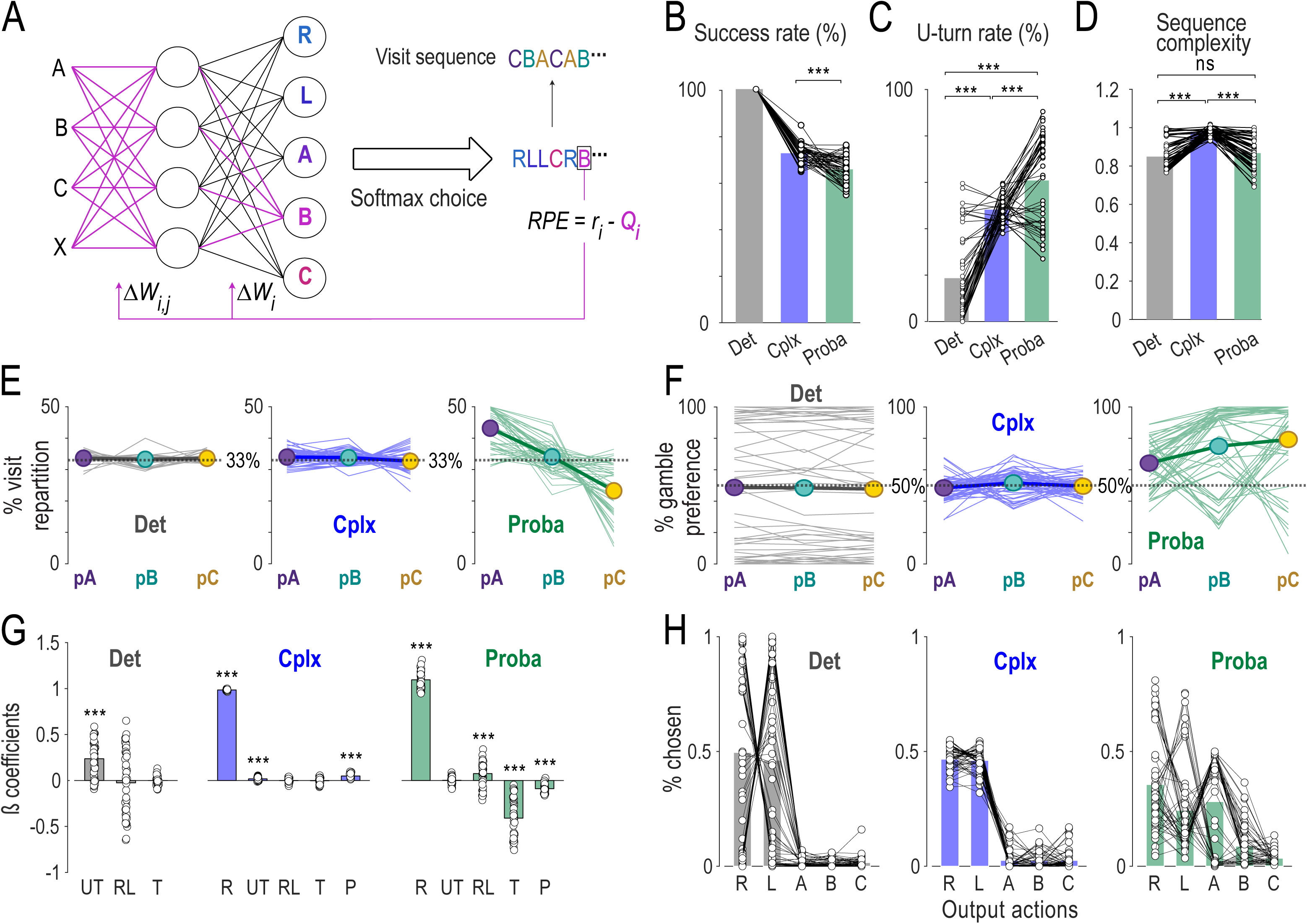
A deep RL network captures behavior and DA signals. **A.** Schematic of the deep RL network, learning rule and sequence generation. The network outputs the values of a set of five actions: R: right; L: left; A, B and C: going to point A, B and C respectively. **B-D.** Comparison across rules of **B.** success rate, **C.** sequence complexity and **D.** Uturn rate **E.** Proportion of target visits **F.** Choice preference at each gamble. The latter shows a bias for circular foraging in Det, randomness in *Cplx* and exploitation in *Proba*. For Det and Cplx, the figure displays the preference for going right at each location, and for proba it displays the preference for the highest-value option at each location. **G.** Regression coefficients of a GLM at the end of Det, Cplx and Proba. Abreviations stands for: R: reward/omission; UT: forward/uturn; RL: right/left; T:target; P: previous trial’s reward/omission. **H**: % of chosen action among the five output of the network. Data are shown as individual points for each simulation, and the bars represent the mean. N=50 simulations.

When exposed sequentially to the Deterministic, Complexity and Probabilistic rules, the deep RL agent reproduced (**Fig. 6B–F**) the main behavioral signatures observed in mice (**Fig. 2C–G**). Success rate decreased and U-turn rate increased at the Det to Cplx transition and at the Cplx to Proba transition (**Fig. 6B–C**). Sequence complexity increased specifically under Cplx (**Fig. 6D**), and the distribution of target visits and choice orientation reflected circular foraging in Det, random-like exploration in Cplx, and exploitation of high-probability targets in Proba (**Fig. 6E–F**). Thus, a single agent, with fixed architecture and learning rules, recapitulated the diversity of strategies expressed by animals across contexts.

To compare the model’s internal “DA” signal with experimental data, we treated the trial-by-trial RPE as a proxy for NAc DA transients and applied the same GLM analysis to model-generated RPEs as to the recorded DA signals (**Fig. 6G**). The dominant predictors of model RPEs shifted across rules in close agreement with the data: direction in Det, current and previous outcomes in Cplx, and target identity (reward probability) in Proba (**Fig. 6G)**, with only a minor contribution of recent history in Proba. Thus, without explicitly encoding rule identity, the network learned input–action mappings that induced rule-specific RPE modulations aligned with the representational structure inferred from NAc DA.

Inspection of the network’s output usage (**Fig. 6H**) clarified how these representations emerged. In Det, the agent relied predominantly on a single motor output (Left or Right), implementing a circular policy (clockwise or counter-clockwise) with few U-turns. In Cplx, Left and Right values converged toward similar levels and were used more evenly, generating high sequence variability and approximating a value-independent, randomness-seeking strategy. In Proba, the agent combined spatial and motor outputs to preferentially visit high-probability targets, yielding stable spatial preferences and frequent U-turns (**Fig. 6H**). Consistent with these learned value landscapes, Det produced a directional asymmetry in RPEs (smaller RPEs along the dominant direction than for U-turns), Cplx produced stronger positive RPEs after prior failures, and Proba produced target-dependent RPEs tracking exploitation of p100 and p50.

Together, these results show that a simple deep RL agent trained on the three rules can (i) reproduce the rule-specific behavioral policies observed in mice and (ii) generate RPE signals whose GLM fingerprints match rule-dependent NAc DA modulations, supporting the idea that flexible mappings from task features to RPEs can arise naturally from representation learning. Building on this, we next asked how these DA-coded representations relate to the *time-resolved* adaptation of choice policies across sessions in both animals and model.

### DA representations track behavioral adaptation across rules

We next asked how the evolution of decision policies across sessions relates to changes in DA dynamics, and whether this coupling is consistent with the deep RL framework. We assessed the evolution of the betas from the GLM analysis of DA variations across sessions, along with the evolution of the betas from the same analysis performed on the RPE generated by the deep RL model (see **Fig. S9A-C** for the evolution of behavior across session). By construction, these beta coefficients capture how strongly the internal value representation differentiates between options or outcomes, and can thus be directly related to evolution of choice bias and success rate.

In the Deterministic rule, the biased circularity profile could emerge from a value discrimination between two directional actions (right vs left), as suggest the evolution of βRL across sessions and its tendency to correlate with the circularity profile of mice (**Fig. 7A–B**). This behavior-RPE structure is strongly predicted by the deep RL model (**Fig. 7C–D**).

**Figure 7:**
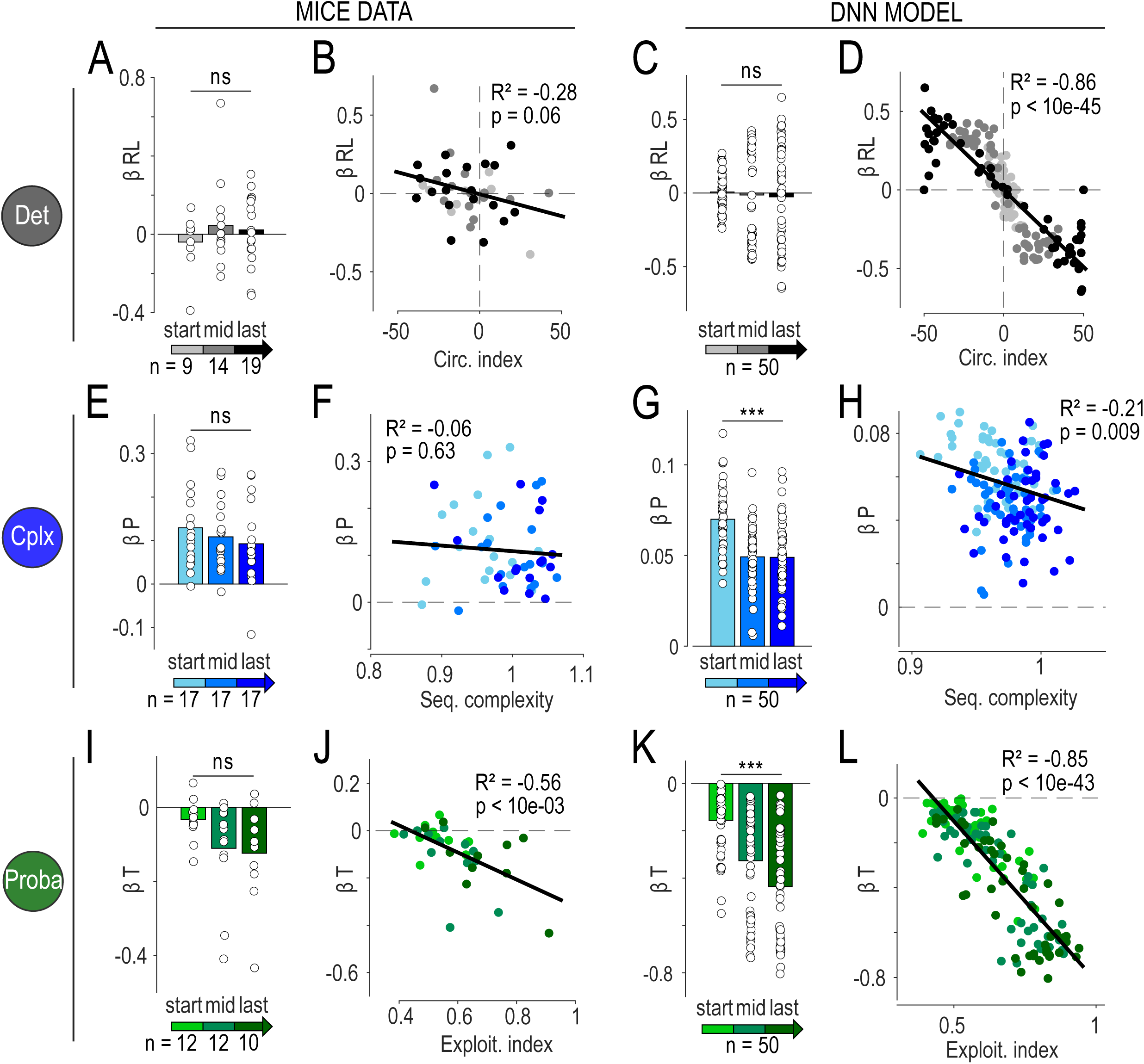
Deep neural network shows similar performance and learning trajectories than mice behavior. **A, E, I.** Evolution of dependence of DA RPEs to principal predictor variable within each rule: **A.** ‘right/left’ (RL) in Det, **E.** previous outcome (P) in Cplx, **I.** ‘target’ (T) in proba. The evolution is shown across sessions spent in each rule with division of sessions into three categories ‘start’, ‘mid’ and ‘last’ which correspond to different stages in the training schedule. **C, G and K** show the same analysis performed on the deep RL simulations data. In figures **B,D,F,H,J,L** each data point represent a mouse (**B,F,J**) or a deep RL simulation (**D,H,L**) at one of the three stages of training schedule. **B, D.** In Det, linear regression between circularity index and dependence of RPEs to ‘right/left’ (RL) for mice (**B**) and deep RL simulation (**D**). **F, G.** In Cplx, linear regression between sequence complexity and dependence of RPEs to ‘previous outcome’ (P), for mice (**F**) and deep RL simulation (**G**). **J, L.** In Proba, linear regression between exploitation index and dependence of RPEs to ‘target’ (T) for mice (**J**) and deep RL simulation (**L**).

Upon transition to the Complexity rule, both mice and the deep RL agent initially experienced high omission rates because the previously optimal circular policy produced low sequence variability (**Fig. S9B**). Mice improved performance by increasing U-turns and redistributing them across trials to generate more variable sequences. Although U-turns could locally increase reward probability (**Fig. S10A**), animals did not adopt a simple “U-turn after omission” heuristic; instead, performance improved through a more global reorganization of sequences (**Fig. S10B–D**). The deep RL agent progressively equilibrate the value of the competing actions R and L, inducing a high entropy policy. As for the mice, this results in a redistribution of U-turns across sequences and a progressive increase of success rate. Notably, the emergence of a high-entropy policy did not require changes in the softmax exploration temperature β, which was held fixed. DA variations remained dominated by recent outcome history (larger responses after a prior reward than after a prior omission), with a tendency of this effect to decrease across sessions, and no correlation with improvements in sequence complexity or success rate (**Fig. 7E–F, Fig. S10E–F**). Regarding the deep RL agent: the history dependent RPE is already present since the start of complexity and decrease across sessions (**Fig. 7G–H**). The model provides a network-level mechanism for this effect with the presence of the hidden-layer which makes learning not confined to the selected output: updates propagate through common units and can partially influence other actions. Deeper simulation analysis demonstrate that this property could not be explained with a 1-layer model (**Fig S11**). Regarding the correlation between history effect and complexity of sequence of the deep RL agent (**Fig 7H**), it results from progressive evolution of both variable across sessions and not from a causal relationship. Thus, in this context, DA transients reflect a history-dependent RPE based on an abstract contingency (sequence variability), while policy adaptation proceeds through reconfiguration of representations and values rather than through DA-driven tuning of the softmax exploration temperature β.

The situation changed in the Probabilistic rule. Mice again exhibited an initial drop-in reward rate, now because targets differed in reward probability and the previously acquired variable policy was not yet aligned with these probabilities (**Fig. S9C**). Across sessions, animals increased success rate by biasing choices toward higher-probability targets, at the cost of more U-turns and more stereotyped trajectories.

In parallel, dependance of DA signal to target identity grew over time, and sessions with stronger sensitivity of DA to target identity exhibited stronger preference for high-probability targets (**Fig. 7I–J**). The deep RL agent mirrored these dynamics: under Proba, learned values diverged across targets, producing increased RPE sensitivity to target identity and a progressive shift toward exploitation (**Fig. 7K–L**). Across both Cplx and Proba, β remained constant, indicating that apparent changes in “exploration/exploitation tradeoff” can arise from changes in learned value landscapes and internal representations rather than from explicit modulation of a global exploration parameter (β).

## Discussion

By recording dopamine (DA) release in the nucleus accumbens (NAc) while mice adapted to multiple reward rules in a single spatial task, we asked how a canonical reinforcement-learning variable—reward prediction error (RPE)—is implemented when the relevant state space itself must change. Across three reward contexts, we found that DA transients did not simply encode a fixed scalar “value error” tied to physical actions or locations. Instead, DA responses aligned with different internal descriptions of behavior in each rule: direction-based in the Deterministic context, history-based in the Complexity context, and probability-based in the Probabilistic context. By combining GLM-based model comparison, classical RL models with hand-specified state features, and a deep RL agent that learns its own internal representations, we show that NAc DA behaves as an RPE computed over flexible, learned action representations, rather than over a rigid, experimenter-defined state space.

First, our results confirm and extend a consistent pattern observed across the dopamine literature, wherein phasic DA carries information regarding both the obtained value and the RPE upon delivery or omission of an expected reward ^8,13–22^. It remains unclear whether this response stems from direct stimulation of MFB DA fibers, resulting in DA release in the NAc, or whether it reflects a subjective value mediated by circuits beyond the DA system alone ^37,38^. While ICSS evoked positive transients across contexts, their amplitude was strongly modulated by expectation. Unexpected rewards and omissions triggered positive and negative transients respectively, consistent with classical RPE encoding ^8,9,22,32^. Thus, our data reinforce the view that mesolimbic DA provides a relatively low-dimensional teaching signal that can be read as a real-time RPE, even in freely moving animals performing spatially extended tasks ^8,22,27,39,40^. At the same time, the rule-dependent structure of DA modulation shows that this RPE is not defined over a fixed “trial” or “arm” variable, but over an internal representation that can be reconfigured as the task demands change.

Second, mice demonstrated substantial flexibility not only in their policy but also in the internal task-state/action representation used for value learning. In the Deterministic rule, optimal performance was achieved by adopting a stereotyped circular policy with few U-turns, and NAc DA was best explained by an RPE defined over movement direction (U-turn vs forward), consistent with extensive evidence that phasic dopamine in the nucleus accumbens reports reward prediction errors for well-learned action-outcome contingencies^41^. In the Complexity rule, by contrast, the same circular policy became detrimental because it produced low sequence variability. Mice recovered reward rate by increasing U-turns and distributing them across trials, thereby raising global sequence complexity^10^. This aligns with operant-learning literature showing that behavioral variability can itself be shaped by reinforcement contingencies and brought under contextual control^42,43^. Strikingly, in this rule DA no longer differentiated directions or targets, but instead was best accounted for by an RPE defined over recent outcome history, consistent with an abstract contingency^33,44^ that rewarded variability. Finally, in the Probabilistic rule, animals exploited fixed reward probabilities associated with locations: they biased choices towards high-probability targets, accepting more U-turns and more stereotypy. In this context, NAc DA reflected an RPE over target identity and reward probability, with dependence of DA RPEs to locations growing in parallel with the behavioral preference for high-probability sites, and consistence with classic work showing that dopamine signals incorporate reward probability/uncertainty and subjective value^8,9,25,26^. Together, these observations indicate that mice not only adjust policy, but also switch which features of actions and outcomes are treated as the relevant “state” variables for prediction and error computation, a core idea in task-state/representation learning frameworks ^1,6,45^, and that NAc dopamine tracks these representation shifts, consistent with the view that RPE-like teaching signals can accompany (and potentially drive) adaptive reconfiguration of learned representations^46^.

Beyond establishing a rule-dependent mapping between task features and DA-like RPEs, the deep RL agent offers a minimal mechanistic account of *how* such mappings can emerge when actions admit partially redundant descriptions. In our architecture, motor (Left/Right) and spatial (“go to A/B/C”) outputs can lead to the same choice, creating a credit-assignment problem in which the agent must attribute outcomes to one of several partially overlapping action descriptions^47,48^. This simple “credit assignment under redundancy” naturally produces the stereotyped circular policy in the Deterministic rule: value concentrates on one directional output, making alternative choices effectively low-value deviations, and generating the directional RPE asymmetry that matches the DA U-turn effect. Crucially, the same architecture can also reproduce the Complexity signature—where DA is best captured by a history-dependent, trial-level representation—because learning is not confined to the chosen output. Weight updates propagate through shared hidden units, so that updating the value of a selected action also partially updates the values of other, unchosen actions that rely on overlapping hidden-layer features. As a result, the hidden layer rapidly comes to encode a coarse estimate of recent reward rate (or global success), which is then broadcasted to multiple outputs. This provides a natural explanation for why the “previous omission lead to larger subsequent reward transient” effect is present from the beginning of the Complexity rule, both in the model and in DA recordings, without requiring a slow, rule-specific accumulation process. More broadly, these results support a view in which dopaminergic teaching signals are computed over *learned* internal representations rather than fixed experimenter-defined state spaces. In the deep RL agent, plasticity in hidden layers both constructs such representations and shapes the ensuing RPE signal, suggesting a plausible biological mechanism: the same dopaminergic error signal could, via synaptic plasticity at multiple stages, progressively tune upstream networks to represent precisely those features that best predict reward in a given context^47^. An important implication is that individual “profiles” within a rule may emerge downstream of this shared rule-level representation, through biased readout and attribution over the learned action manifold—an idea that can be directly tested by quantifying whether hidden-layer activity is more strongly structured by rule identity than by within-rule phenotypes.

The joint analysis of the evolution of dependence of DA RPEs of mice and RPE of the model to predictors across sessions further constrains how we should think about DA’s role in exploration and policy adaptation. In the Complexity rule, mice increased sequence variability and improved performance without developing any systematic DA dependence to specific actions, but displayed DA dependence to recent reward vs omission. The deep RL agent exhibited the same qualitative pattern: it recovered reward rate by making repetitive sequences predict low return and by increasing the entropy of its policy, yet did so without any change in the softmax temperature β, which was held constant across rules. This implies that the apparent increase in “exploration” in Cplx can emerge from changes in representations and value estimates, rather than from a dedicated DA-driven modulation of a global exploration parameter. Conversely, in the Probabilistic rule, both mice and model showed a tight coupling between the growth of DA-RPE dependence to targets and the consolidation of an exploitative policy favoring high-probability locations. Here, value-based asymmetries in DA (or RPE) and policy refinement evolved in parallel, again under a fixed β in the model, because the learned value landscape became more uneven. These comparisons argue that at least part of the DA–behavior relationship in our data can be explained by standard RPE-based value learning over flexible state representations, without invoking an explicit DA control signal for exploration temperature. This does not preclude additional, slower or circuit-specific mechanisms for tuning exploration–exploitation balance^34,49,50^, but it suggests that they are not required to account for the pattern we observe.

Several limitations of our approach point to directions for future work. First, fiber photometry in NAc measures a population-averaged DA signal and cannot resolve heterogeneity across projection-defined subpopulations or across core vs shell. Different subsets of DA inputs may carry distinct RPEs or emphasize different state features^23,51,52^, which our bulk signal effectively averages. Second, our task manipulates reward structure in a relatively simple, discrete set of rules; in more naturalistic environments^8,53^, changes in contingencies may be more gradual or partially observable, and representation learning may involve hierarchical or belief-state-like structures that go beyond the models tested here. Third, our deep RL agent is intentionally minimal and does not include mechanisms such as meta-learning of learning rates, explicit uncertainty monitoring, or interactions with other neuromodulatory systems. Extending the framework to tasks with richer temporal structure or partial observability, and to models in which DA interacts with noradrenergic or cholinergic signals, will be essential to understand how different neuromodulators jointly shape internal models and exploration strategies.

In conclusion, our findings reveal that dopaminergic signals in the NAc carry a robust RPE-like teaching signal, but that this error is anchored to an internal, learned description of action and context that changes across reward rules. Behavioral adaptation in our task involves both policy changes and representation changes, and NAc DA tracks these representation changes in a way that can be captured by a single deep RL architecture. These results support a view of DA as a neural marker of representation learning in complex environments: not merely reporting errors in a fixed state space, but reporting errors in whatever task representation the brain has currently constructed to solve the problem at hand.

## Acknowledgements

This work was supported by the Centre National de la Recherche Scientifique CNRS UMR 8246 and 8249, INSERM U1130. We are grateful to the animal facilities (IBPS Sorbonne University) and Otilia de Oliveira and Emilie Tubeuf at ESPCI animal facilities. We also thank Jérémie Naudé and Clément Solié for their comments on the manuscript. We thank Deniz Dalkara and the viral core facility at the Vision Institute (Paris) for production of the AAVs. This work has received support under the Major Research Program of PSL Research University “PSL-Neuro” launched by PSL Research University and implemented by ANR (ANR-10-IDEX-0001). Our team is affiliated to DIM C-BRAINS, funded by the Conseil régional d’Ile-de-France.

## Fundings

The Foundation for Medical Research (FRM, Equipe FRM DEQ2013326488 to P.F, PhD fellowhsip ECO201806006688 to J.J., Fourth-year PhD fellowship FDT201904008060 to MC), the French National Cancer Institute Grant TABAC-19-020 and SPA-21-002 (to P.F.). French state funds managed by the ANR (ANR-19-CE16-0028 Bavar to PF, ANR-23-CE37-0017 VarSeek to PF). Memolife Labex, fourth-year PhD fellowship (to JJ and EV).

## Authors contributions

Conceptualization: MC, PF

Injection and implantation surgeries: MC

Behavioral experiments: MC, AG, LK

Fiber photometry recordings: MC, AG

Intracardiac perfusions and immunohistochemistry: MC, AG, EV, TLB

Data analysis: MC, AL, PF

Modelling: MC, AL, PF

Deep RL: AL, PF

Setups development: MC, JJ, AM, EB, SD, PF

Funding acquisition: PF

Writing - original draft: MC, AL, PF

Writing - review and editing: MC, TLB, AM, PF

## Competing interests

Authors declare that they have no competing interests

## Materiel and Methods

### EXPERIMENTAL MODEL AND SUBJECT DETAILS

Experiments were performed on adult C57Bl/6Rj wild-type mice (Janvier Labs, France). Both male and female mice, weighing 20-30 g and 8 weeks old at the time of surgery, were used for behavioral experiments. Only male mice were used in the GRAB_DA_ fiber photometry cohorts. For cre-dependent GCaMP experiments, DATiCre male mice were used. All mice were kept in an animal facility where temperature (20 ± 2°C) and humidity were automatically monitored and a circadian 12/12h light–dark cycle was maintained. All experiments were performed in accordance with the recommendations for animal experiments issued by the European Commission directives 219/1990, 220/1990, and 2010/63, and approved by Sorbonne University and ESPCI.

### METHOD DETAILS

#### AAV production

AAVs for GRAB_DA2m_ (pXR1-AAV-hSyn-GRAB-DA4.4) were produced as previously described ^54^ (using the cotransfection method from plasmids generously provided by Dr. Yulong Li ^36,55^ and purified by iodixanol gradient ultracentrifugation^56^). AAV vector stocks were tittered by quantitative PCR (qPCR) ^57^ using SYBR Green (Thermo Fischer Scientific). AAV vectors for GCaMP6f (AAV1-EF1a-DIO-GCaMP6f-P2A-nls-dTomato) and GCaMP7c (pGP-AAV1-syn-FLEX-jGCaMP7c variant 1513-WPRE) were directly ordered from Addgene.

#### Intracranial self-stimulation (ICSS) electrode implantation

Male and female WT mice were anaesthetized with a gas mixture of oxygen (1 L/min) and 1-3% of isoflurane (Piramal Healthcare, UK) and then placed into a stereotaxic frame (Kopf Instruments, CA, USA). After the administration of a local anesthetic (Lurocain, 0.1 mL at 0.67 mg/kg), a median incision revealed the skull, which was drilled at the level of the median forebrain bundle (MFB). For ICSS, a bipolar stimulating electrode (PlasticOne 2 channels, stainless steel, 10 mm) was then implanted unilaterally (left or right, randomized) in the brain using the following stereotaxic coordinates (from bregma according to Paxinos atlas): AP −1.4 mm, ML ±1.2 mm, DV −4.8 mm from the brain). Dental cement (SuperBond, Sun Medical) was used to fix the implant to the skull. An analgesic solution of buprenorphine at 0.015 mg/L (0.1 mL/10 g) was delivered prior to awakening from the surgery and, if necessary, the following recovering days. After stitching, mice were placed back in their home-cage and had a minimum of 5 days to recover from surgery. The efficacy of electrical stimulation was verified through the rate of conditioning during the deterministic setting (see Intracranial Self Stimulation (ICSS) bandit task). Out of the 54 mice implanted (27 for each sex), 49 were included in the results (23 males and 26 females).

#### Virus injections and fiber photometry recordings

3 cohorts of WT male mice (total of 24) were anaesthetized (Oxygen 1 L/min, Isoflurane 1–3%) and implanted with an ICSS electrode as described above. They were then injected unilaterally (randomized left/right side and ipsi/contralateral side regarding the ICSS electrode) in the NAc lateral shell (1 µL, coordinates from bregma: AP +1.45mm; ML ±1.55mm; DV −4.05mm from the skull) with an adeno-associated virus ^36,55^ to express GRAB_DA2m_. An optical fiber (200 µm core, NA = 0.39, Thor Labs) coupled to a metallic ferule (1.25 mm) was implanted 100 µm above the injection site in target region and cemented to the skull with blackened cement. 5 DATiCre male mice followed the same procedures for GCaMP experiments in the VTA (1 µL, coordinates from bregma: AP - 3.10mm; ML ±0.50mm; DV −4.20mm from the brain), 3 of them with GCaMP7c and 2 with GCaMP6f. Viral expression typically took 10-15 days to achieve a satisfying signal and lasted for up to 3 months. However, some mice exhibited a shorter duration of expression and were therefore excluded for the analysis of later sessions. Although the mice performed the task on a daily basis, fluorescent recordings were made only every 2 or 3 days to prevent sensor bleaching. Low power (100-200 mA) LEDs (465 nm and 405 nm, Doric Lenses) coupled to a patch cord (500 µm core, NA = 0.5, Prizmatix) were used for optical stimulation of the sensors in lock-in mode (572.205 Hz for the 465 nm LED, 208.616 Hz for the 405 nm LED) and collection of 520 nm fluorescence. 405 nm was used as the isobestic wavelength. The optical stimulation patch cord was plugged onto the ferrule during all experimental sessions, even those without recordings, to habituate animals and control for latent experimental effects. After the daily session, a short recording of the autofluorescence signal *F(auto)*, coming from the patchcord only, was performed with same LED intensities, no animal plugged and room in the dark. Raw 520 nm fluorescence was demodulated by the software (Doric Lenses) to extract 465 nm and 405 nm signals. The 405 nm signal was visually checked to account for instability artefacts coming from head movements or patch cord unplugging during the session, and if needed correct the associated 465 nm signal accordingly, otherwise it was not used for signal treatment. 465 nm signal *F_i_* follows several treatment steps according to this formula:

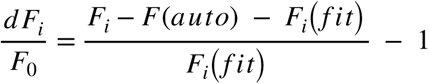

First *F_i_* is subtracted with the constant value of autofluorescence *F(auto)* measured with patch cord only, improving drastically the signal-to-noise ratio. Then, largest transients induced by ICSS were excluded to perform a smoothing on the subsequent truncated signal. We then computed a mono-exponential fit *F_i_*(*fit*) on this smoothed signal, which was also subtracted to *F_i_* at each time point *i* to account for exponential decay. The result is then divided by the same *F_i_*(*fit*) at each time point *i* to normalize the signal around 1, and subtracted by the constant 1 to normalize to 0 and obtain positive or negative transients as dF_i_/F_0_ over an entire session (5 or 10min). In order to aggregate signals coming from different sessions for each mouse, and then pool mice for the analysis, we also applied a z-scoring on dF_i_/F_0_ over each entire session.

#### Intracranial self-stimulation (ICSS) bandit task

The ICSS bandit task ^9,10,34,35^, took place in a circular open-field with a diameter of 68 cm. Three explicit square-shaped marks (2 × 2 cm) were taped in the open field, forming an equilateral triangle (side = 35 cm). Entry in the circular zones (diameter = 6 cm) around each mark was associated with the delivery of a rewarding ICSS stimulation. A LabVIEW (National Instruments) application precisely tracked and recorded the animal’s position with a camera (20 frames/s). When a mouse was detected in one of the circular rewarding zones, a TTL signal was sent to the electrical stimulator, which generated a 200 ms train of 5 ms biphasic square waves pulsed at 100 Hz (20 pulses per train). Two consecutive rewards could not be delivered on the same target, which motivated mice to alternate between targets and therefore generate sequences of binary choices. ICSS intensity was adjusted, within a range of 15-200 µA, during early conditioning sessions, so that mice would achieve between 50 and 120 visits per session (5 min duration) for two successive sessions. ICSS intensity was then kept constant for all the experiments, even when reward delivery rules changed. Mice with insufficient scores were excluded. Different reward delivery rules were used, and all animals went through all three protocols successively. The first is a deterministic (Det) setting, with 10 to 17 daily sessions of 5 min. All zones were associated with an ICSS delivery (P = 100%). The second, described previously in ^10^, is a complex (Cplx) setting where a grammatical complexity algorithm ^58^ analyses online the choice sequence that the mouse is producing, calculates the complexity of two potential sequences of length 10 (9 past targets + next target among the 2 available) and gives a reward only if the complexity of the sequence increases. Repeating patterns of low complexity will therefore lead to series of omissions, while increasing variability will increase success rate. Mice did daily sessions during 15-20 days. The third setting is probabilistic (Proba): each target is associated with a probability to obtain an ICSS stimulation among three (P = 25%, P = 50%, P = 100%), as described previously ^9,34,35^. The probabilities at each location were pseudo-randomly assigned per mouse, and 15-20 sessions were performed. 2 cohorts of both male and female mice followed deterministic, complexity and probability settings successively, with no fluorescent sensor expression. Three cohorts of male mice expressing GRAB_DA_ and implanted with an optical fiber implantation followed different settings: *i)* the first cohort performed only Det and Cplx, and recordings started only at the end of Det, *ii)* the second and third cohorts performed Det, Cplx and Proba, with recordings starting at the beginning of Det, and performed also some control experiments (especially, unexpected rest cage and off-target ICSS). Consequently, there is variation in animal numbers among conditions in the figures. Finally, one cohort of DATiCre male mice was tested in Det and Cplx only. We defined three analysis windows within each condition (see Fig 7) - Start, Late, and End - corresponding to the following experimental session ranges: Deterministic (Det) sessions 6–7, 11–12, and 14–17; Complexity (Cplx) sessions 1–4, 5–8, and 10–13; and Probabilistic (Proba) sessions 1–2, 3–6, and 11–16.

### QUANTIFICATION AND STATISTICAL ANALYSIS

#### Behavioral measures

For all those groups, the following measures were analyzed with custom codes in Python (using mostly Numpy and Pandas libraries, on PyCharm CE) and compared throughout the different rules: *i)* number of visits, *ii)* success rate, *iii)* time-to-goal, *iv)* choice repartition (proportion of visits at each location), *v)* percentage of U-turn (target n = target n+2) and *vi)* sequence complexity (applying the same complexity algorithm calculation but offline and for all choices during a session). Furthermore, the ICSS bandit task can be seen as a Markovian decision process: every transition can be considered as a binary choice between two options, since a zone cannot be reinforced twice in a row. The sequence of choices per session results from the succession of three specific binary choices, or gambles. For deterministic and complexity, G_C_ = P(A|C) would be the total number of visits in target A divided by the total number of visits in targets A and B, when the animal is in target C. Similarly, G_A_ = P(B|A) and G_B_ = P(C|B). A gamble above 50% indicates that the animal has a preference for moving clockwise (or below 50% for moving counter-clockwise). In probabilistic, direction of conditional probabilities does not follow spatial repartition of locations, but rather preference for the high value option: G_25_ = 100% vs 50%, G_100_ = 50% vs 25% and G_50_ = 100% vs 25%. Applying this principle at each choice, those 3 gambles can be aggregated into single values to give circularity index (going in circle, no matter clockwise or counter-clockwise), exploitation index (always preferring the highest value option) or repetition index (always making the same choice at given gamble, no matter the direction or exploitation).

#### Fiber photometry analysis

All treatments and analyses were performed in Python using custom codes (mostly Numpy and Pandas libraries). After cleaning and processing each session signal to obtain dF/F values and z-scored dF/F values, events of interest were extracted to align the signal in [-3s:3s] time window in dataframes, t_0_ being the exact time of location entry (triggering reward delivery or omission), with 1kHz sampling. Session-wise averages of given conditions for each mouse were then extracted, and averaged again over multiple mice for statistical analyses. In some conditions, especially when events of interest were rare (some scenarios of rewards or omissions chains in complexity, or some scenarios of locations transition in probabilistic), two or more sessions from one animal were pooled as if they were one (for instance, the last two sessions in a given context) to have enough trials for each animal in this condition. For the same reason, the third cohort of GRAB_DA_ mice followed 10 min long sessions (instead of 5 min) in Cplx and Proba settings, with no particular effect on the overall quality of the signal, nor the duration of GRAB_DA_ expression (up to 3 months). For GRAB_DA_, rewards-elicited positive transients typically peaked around 250 ms after location entry (duration of ICSS being 200 ms) and decayed during a bit less than 1s: we therefore extracted maximum and mean of the signal in a 1 s window post location entry. Omissions-elicited negative transients were longer, reaching their minimum around 800 ms after location entry and taking roughly 700-800 ms to go back to baseline: we therefore extracted minimum and mean of the signal in 1.5 s window post location entry. For GCaMP, kinetics depended on the sensor used: peaks reached maximum value around 250-300 ms post location entry for GCaMP6f and 350-400 ms for GCaMP7c, while dips reached minimum value around the same time (900-1050 ms post location entry) for both sensors. However, return to baseline after reward-induced peaks was much shorter for GCaMP6f (500-600 ms post location entry) than for GCaMP7c (2-3 s). For some correlation analyses (using SciKit Learn Python library), especially the ones regarding z-scored peaks or dips amplitude regarding outcome chain history, all trials of all mice were pooled together in a given condition.

#### Generalised Linear Model (GLM) approach

GLM was performed in Python using custom codes (StatsModels or SciKit Learn library). To disentangle multiple factors that could explain DA signal, due to high degree of behavioral and task-related variables correlated to each other from one trial to the next, we designed a generalized linear model where a variable *Y* is explained by a linear combination of multiples variables *X_i_*, each of them weighted by a parameter *w_i_*, plus a residual (or intercept) *w*_0_.

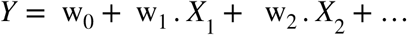

The model aims at fitting variations of *Y* by determining the weights *w_i_* and their significance. Dependent variable *Y* was post location entry 1s average for reward-induced peaks or 1.5s average for omission-induced dips. Multiple *X_i_* variables have been used, namely: *i)* reward or omission at previous and current location, *ii)* Forward or U-turn at previous trial, *iii)* current target visited (spatially A, B or C, or in Proba p_100_, p_50_ or p_25_), and *iv)* time since last stimulation (in Restcage stimulation condition). A single GLM was applied for each mouse in a given condition, then ^w^*i* parameters resulting from all those GLMs were averaged among mice, and the average was statistically compared to 0. Significance, either with positive or negative weight, indicates that this variable explains part of DA variations.

#### Reinforcement Learning (RL) models

We used Reinforcement Learning (RL) to compute Reward Prediction Errors (RPEs) from actual mice choice sequences and see how they match DA data. Before each trial, the agent contains a set of expected values for each possible action. As one of these actions is selected, it leads to either a reward (V_obtained_ = 1) or an omission (V_obtained_ = 0), then RPE is calculated as V_obtained_ - V_expected_, and a new expected value of this action is fed back into the agent’s set for next trials. From both behavioural and photometry results, we hypothesised and tested three possible value representations in the bandit task. First, we proposed a simple, one-order representation “going to any target” or “performing any trial” to get a reward. In this case, all trials are similar, regardless of target or trajectory choices, and we simply compute and update V_expected_ = { V_Any_ } at each trial. Second, a representation of internal directionality with a set of two actions and V_expected_ = { V_Fwd_; V_Uturn_ }. In this case, RPEs are specific and computed separately for each of the two actions. Third, a spatial representation “going to target X” with a set of three actions and V_expected_ = { V_pA_; V_pB_; V_pC_ }. Again, RPEs are computed for each target independently. Modelling the RPE values resulting from each of those three representations allowed us to compare them and determine which simulation better replicates DA data in each context. Initial V_expected_ were set consistently with behavior in the task. For Det End, they were all set to 0.99. For both Cplx End and Proba End, they were set as mean success rate computed from the two previous sessions. For example, for a given mouse, initial V_Uturn_ to initiate the RL model with choice sequence from sessions 9-10 is the proportion of rewarded Uturn trials from sessions 7-8. Exception is for V_p100_ in Proba End where it was also set to 0.99. We arbitrarily tested several learning rates α = {0.001; 0.01; 0.05; 0.2; 0.4}. Results were consistent with experimental data for α = {0.01; 0.05; 0.2; 0.4}. Smaller α (0.001) led to convergence that was too slow considering mice number of trials provided to models, while larger α made convergence in Det too quick. In Fig 3 and Fig S6, α is set to 0.05. We next assumed that in our recordings, DA = V_obtained_ + RPE, and tested which representation accounted most in the error component using GLMs on top of our RL-computed RPEs (taking as input variables V_obtained_ = {1; 0} for rewards or omissions, and theoretical RPEs computed from Model 1, 2 and 3). Similarly, models were applied for each mouse in a given context, then *w_i_* parameters were averaged among mice for each context, and the average was statistically compared to 0. Significant weight indicates that this variable explains part of DA variations. Finally, we extended this compilation of RL-computed RPE values and GLM to fit RPE weights to DA data across sessions and contexts (Fig 4 and Fig S7). In this case, we started RL models with mice choice sequences in Det Start with all V_expected_ equal to zero (naive agents), computed corresponding RPEs and updated corresponding V_expected_. Consistent with mice progressively learning and updating values across sessions and contexts, the final V_expected_ of a given time-point became the initial V_expected_ of the next time point. For instance, from Det Start to Det Late (all V_expected_ becoming closer to 1, but not at the same speed). Or from Cplx End to Proba Start (V_expected_ of each target therefore starting to diverge). To allow for longitudinal comparisons, we next scaled (z-score) our data (both experimental DA and RL models-computed RPEs) on each time point, applied GLMs on each time point, and then compared the weights *i)* across sessions in a given context and at each transition between contexts, and *ii)* each of them regarding its difference with 0.

### DEEP REINFORCEMENT LEARNING MODEL

#### Three-point task simulation

We trained a two-layer deep neural network to predict the values of different actions inside a simulation of the three-point task. The simulation was designed trial-wise. In each trial, the model was located at one of the three points and had to choose an action to go to any of the two available points. We provided the model a set of 5 minimal actions, each corresponding respectively to go right, go left, and go to point A, B or C. After the network predicting the value of these 5 actions, a softmax function was used to convert the output value vector *y* into probabilities of choice using the following formula:

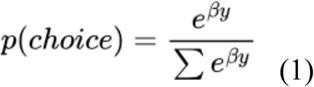

Here β was a choice temperature constant, always set to 9 in our simulations to closely fit mouse data, although other reasonable β values produced qualitatively similar results. The chosen action was then sampled using the probabilities. Note that at each trial, one of the go to A, B or C actions, was made unavailable due to the model being already present at this location. It was thus excluded from the softmax computation. As we gave the model abilities to use location information in the available actions (with A, B, C) we also provided information about its current location in the input. We also gave the model an input that would always be equal to one regardless of the position, to facilitate the non-use of location information if they were not necessary to predict the values. Thus, the three possible input vectors were the following:

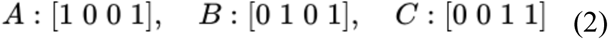

where each corresponds to the location of the model on point A, B or C.

#### Architecture and initialization

The vector of actions values *y* is given by the equation:

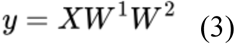

with *X* being the vector of inputs as defined in (2), W^1^ the weights to compute the hidden layer vector from the input vector, and W^2^ the weights to compute the output vector y from the hidden layer vector. We opted for a network with 4 hidden units (same as the number of inputs), even though a different number of hidden units produced similar results (tested with 1 and 10 hidden units). For each simulation, the initialization of W^1^and W^2^ were sampled from a uniform distribution in the interval [-0.01; 0.01] such that initial predicted values from y output vector were close to zero.

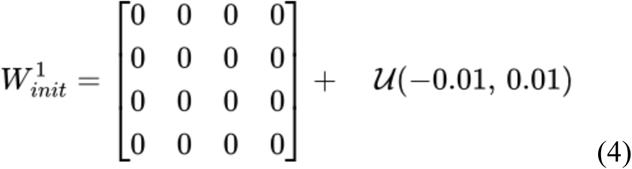

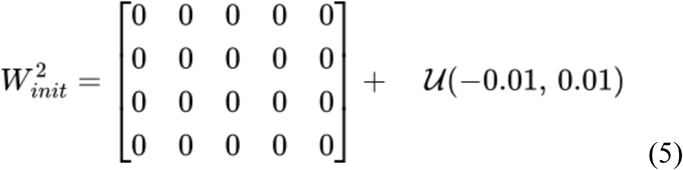

#### Training and Reinforcement

We trained each simulation for 400 trials in each rule set of the three-point task in the same order as the mice: first determinist, then complexity, and finally probabilistic. We updated the weights of the network using stochastic gradient descent at the end of each trial. At each rule switch, the weights of the network were preserved. Each update minimized the following loss function:

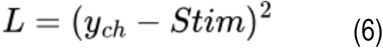

where y_ch_ is the value of the chosen action and Stim. is the result of the choice, taking the value 1 if the model receives a “stimulation” and 0 if not. From this loss, the gradients to update W^1^and W^2^ were computed using chain rule as follow:

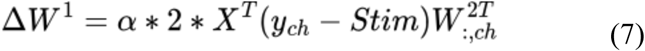

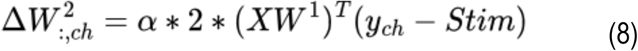

where α is the learning rate and was set to 0.02. Note that the gradient for W^1^were computed by only back-propagating information relative to the computation of the chosen action value, which correspond to the weights contained in only one column of the W^2^ matrix. Similarly, the gradient for W^2^ was a one column vector that would update the weights of the only column of the W^2^ matrix that is necessary to compute the value of the chosen action. Because of this design, the actions were in competition for learning, which could favor the use of different actions depending on the actual rule set.

#### Figures and Statistics

Raw figures were plotted using Python custom codes (mostly MatPlotLib library). Graphics, typography and layout were formatted with Adobe Illustrator. All statistical analyses were computed using Python with Scipy library and custom programs. Results were most frequently plotted as individual data points and mean ± sem. The total number of observations in each group and the statistics used are indicated in figure legends and detailed statistics tables: unless specified, data points indicate the number of mice (N) on which the statistics were performed, and in some cases, they represent number of trials (n) either for one example session from one animal, or from all sessions of all animals in a given condition. Classical comparisons between means were performed using parametric tests (Student’s t-test, or ANOVA for comparing more than two groups, when parameters followed a normal distribution (Shapiro test P > 0.05)), and non-parametric tests when the distribution was skewed (here, Wilcoxon or Mann-Whitney U for one/two samples and whether comparison is paired or not, or Kruskall-Wallis for more than two groups). More complex comparisons with several factors were performed using two-way or mixed ANOVA regardless of normal distribution for simplicity, with no major impact on results interpretation (see Fig S1, sex X session effects). Multiple comparisons were corrected using a sequentially rejective multiple test procedure (Holm). Linear regressions were assessed either with Pearson (parametric) or Spearman (non-parametric) tests. Probability distributions were compared using the Kolmogorov–Smirnov (KS) test. All statistical tests were two-sided. p > 0.05 was considered not to be statistically significant. In some cases, p > but close to 0.05 were indicated in the figure (see Tables of detailed statistics for more information).

#### Fluorescence immunohistochemistry

After completing the successive rules of the task, mice from the GRAB_DA_ cohorts were euthanatized by IP injection of euthasol (0.1mL per 30g at 150mg/kg), immediately followed by paraformaldehyde (PFA) intra-cardiac perfusion, and brains were rapidly removed and post-fixed in 4% PFA for 2 to 4 days. Serial 60µm sections were cut with a vibratome (Leica). Immunohistochemistry was performed as follows: free-floating VTA and NAc brain sections were incubated for 1h at 4°C in a blocking solution of phosphate-buffered saline (PBS) containing 3% bovine serum albumin (BSA, Sigma A4503) and 0.2% Triton X-100, and then incubated overnight at 4 °C with *i)* a mouse anti-tyrosine hydroxylase primary antibody (TH, Sigma, T1299) at 1:500 dilution and *ii)* a chicken anti-eYFP primary antibody (Life technologies Molecular Probes, A- 6455) at 1:1000 dilution, both in PBS containing 1.5% BSA and 0.2% Triton X-100. The following day, sections were rinsed with PBS and then incubated for 3 h at 22–25 °C with *i)* Cy3-conjugated anti-mouse secondary antibody (Jackson ImmunoResearch, 715-165-150) at 1:500 dilution and *ii)* a goat anti-chicken AlexaFluor 488 secondary antibody (711-225-152, Jackson ImmunoResearch) at 1:1000 dilution, both in a solution of 1.5% BSA and 0.2% Triton X-100 in PBS. After three rinses in PBS, slices were wet-mounted using Prolong Gold Antifade Reagent with DAPI (Invitrogen, P36930). Microscopy was carried out with a fluorescent microscope Leica DMR, and images captured in gray level using MetaView software (Universal Imaging Corporation) and colored post-acquisition with ImageJ. Labeling for YFP in the NAc (along with satisfying signal during the task) allowed to confirm GRAB_DA_ expression, and fiber implantation in the NAc lateral shell was also visually checked. Similar procedures were used to check for GCaMP7c and GCaMP6f expression in VTA DA neurons. For GCaMP7c we used the same anti-TH and anti-eYFP antibodies as previously described.

For GCaMP6f we used a sheep anti-TH primary antibody (AB-1542, Milipore) at 1:500 dilution coupled with a donkey anti-sheep secondary antibody (713-165-147, Jackson ImmunoResearch) at 1:500 dilution to highlight DA neurons, and simply used the virus-associated tdTomato to validate expression in the VTA and optic fiber implantation site. For MFB slices, 100 µm sections were performed and slices were directly visualized with visible light to check for ICSS electrode implantations.

#### Statistics and Reproducibility

All experiments were replicated with success (several successive cohorts of mice).

### Supplementary figures

**Fig. S1:**
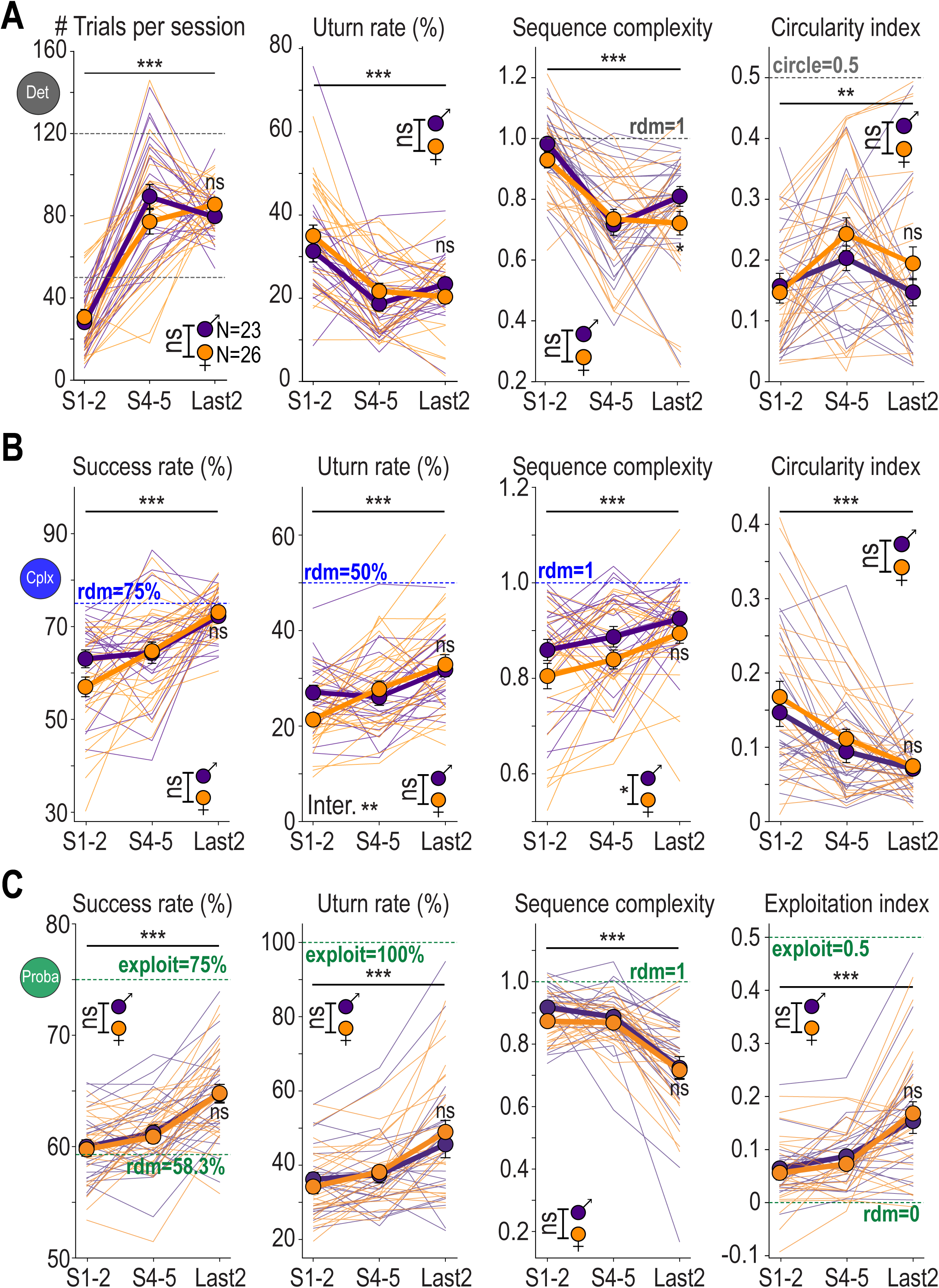
Evolution of decision behaviour across sessions, with no major sex effects. **A. Decision parameters throughout Det sessions for males and females.** Comparison of **(left)** the number of trials per session, **(middle-left)** the Uturn rate, **(middle-right)** the sequence complexity, and **(right)** the circularity index between sessions 1&2, sessions 4&5 and the last 2 sessions in male and female mice. In addition, we also compared the final states (Last2) between males and females. A fully circular mouse would have 0% Uturn, low seq. cplx and 0.5 circul. idx. **B. Same as in A. for Cplx sessions.** A mouse keeping circular strategy would have low success, 0% Uturn, low seq. cplx and 0.5 circul. idx. A random mouse would have 75% success, 50% Uturn, seq cplx = 1 and circul. idx = 0. **C. Same as in A. for Proba sessions.** An exploitative mouse would have 75% success, 100% Uturn, low seq. cplx and 0.5 exploit. idx. A random mouse would have 58.3% success, 50% Uturn, seq cplx = 1 and exploit. idx = 0. (Data are shown as individual points, and mean ±sem. N = 23 male and 26 female mice.)

**Fig. S2:**
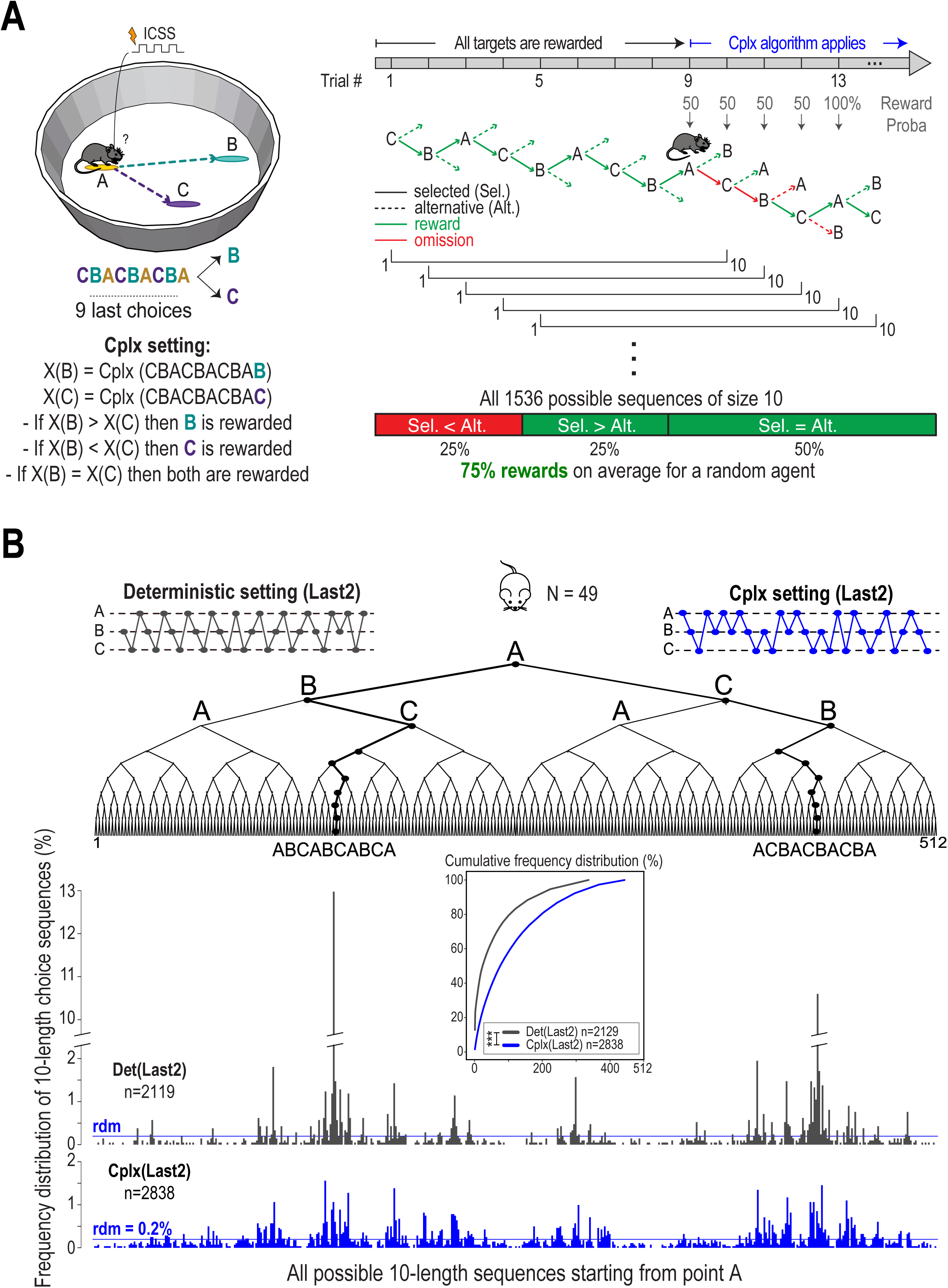
Additional information on the Cplx rule and mice sequence patterns. **A. Detailed schematic representation of the Cplx rule.** The first 9 trials of each session provide deterministic rewards (P=100%) to launch the Cplx algorithm, which then determines at each trial, in a sliding window, which target will lead to a reward by comparing the Lempel-Ziv grammatical complexity of the two potential sequences: 9 past choices + first remaining target VS. 9 past choices + second remaining target. The mouse will be rewarded only if it chooses the target that increases complexity. If both sequences have the same complexity, both targets will be rewarded **(see Methods)**. Taking all possible sequences of size 10 starting from one location, 75% of them are rewarded on the 10^th^ trial. Therefore, a random agent exploring homogeneously this sequences tree will converge to 75% success rate. **B. Distribution of mice choice sequences of length 10 at the end of Det and Cplx.** Two distribution peaks (paths in the decision tree) appear in Det, corresponding to circling behavior (clockwise and counterclockwise), representing together roughly 25% of all produced sequences (among 512 possibilities). In Cplx, these peaks strongly reduce in size, in favor of more distributed visits of all possible sequences. **(Insert)** Cumulative distribution comparison between Det and Cplx (Last2 sessions for each rule). (In B, n is the total number of sequences of length 10, computed from sessions-wise mice successive choices, from N=49 mice both males and females).

**Fig. S3:**
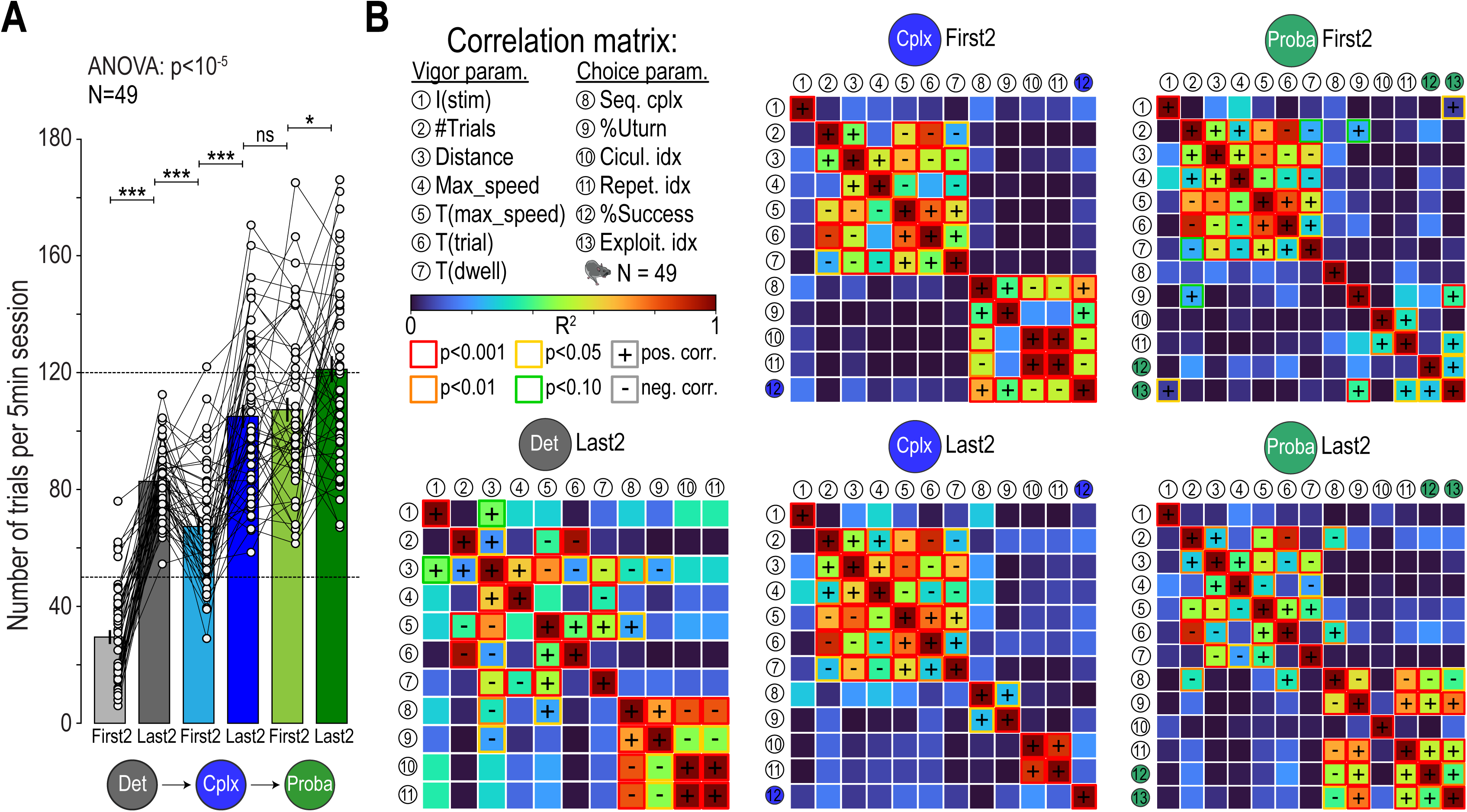
Motivation throughout the task and decoupling between vigor and choice parameters. **A. Evolution of motivation across contexts and sessions.** Comparison of number of trials across contexts and sessions (mean of 2 sessions each time). **B. Correlation matrices between various vigor and choice parameters across mice in different contexts.** Parameters are computed for each mouse as the mean of 2 sessions (either First2 or Last2, for a given context). Each box represents the linear correlation between two parameters (Pearson for parametric, Spearman for non-parametric, each dot being a mouse). The filling color of each box represents the R2 value. The frame color of each box represents the p-value (after Bonferroni correction). The warmer the color, the more those two parameters are significantly correlated. **(Left)** Last2 sessions of Det (11 parameters, x66 Bonferonni correction). **(Middle)** First2 and Last2 sessions of Cplx (12 parameters, x78 Bonferonni correction). **(Right)** First2 and Last2 sessions of Proba (13 parameters, x91 Bonferonni correction). (In A, data are shown as individual points, and mean ±sem. In B, only R2 and corrected p-values are shown with color code. Individual data are available upon request. N is always the number of mice.)

**Fig. S4:**
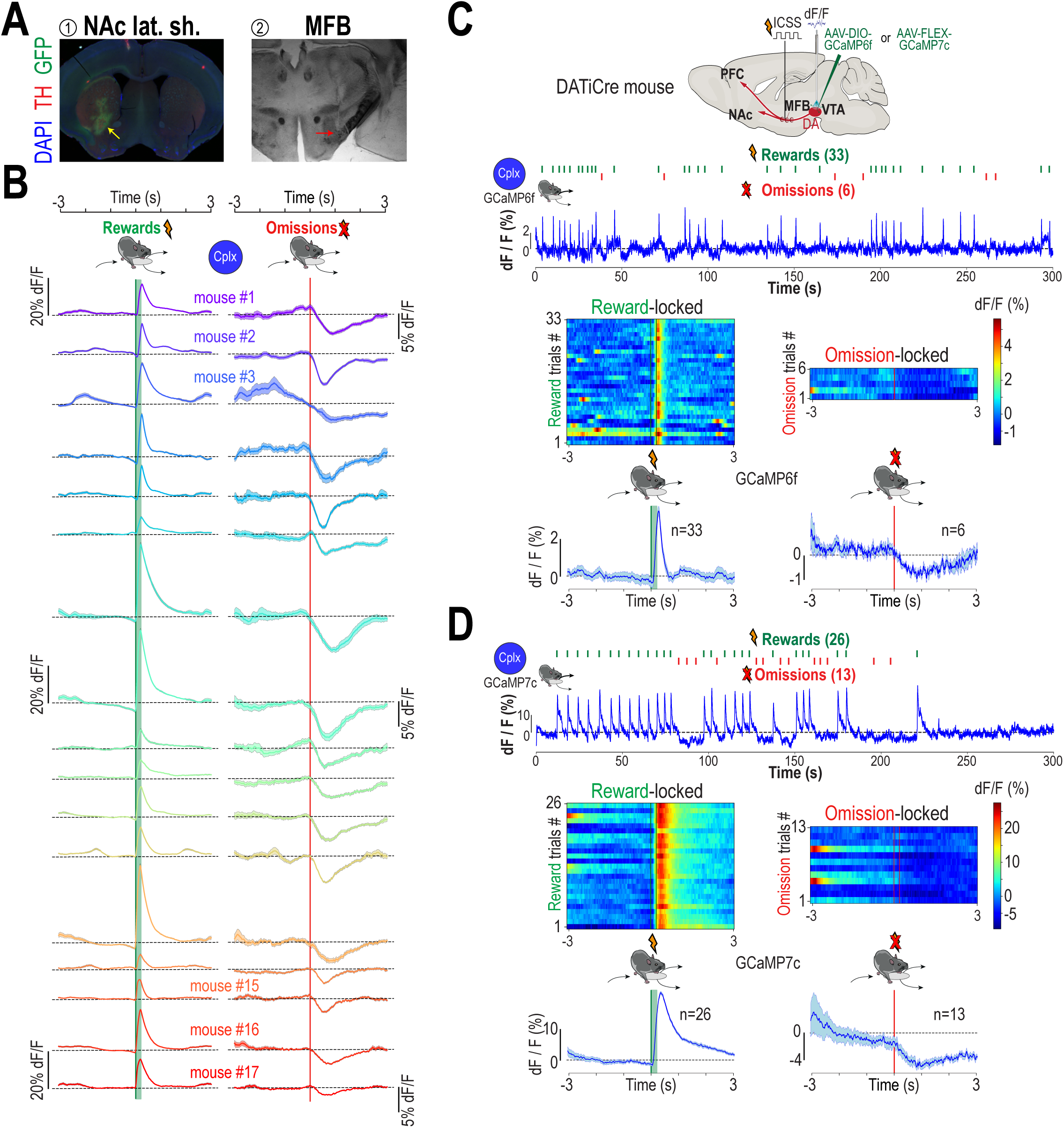
DA fiber photometry signals with Grab_DA_ and GCaMP6f. **A. NAc and MFB slices immunohistochemistry.** Post-hoc verification of optic fiber implant and Grab_DA_ virus expression in the NAc lateral shell (left), and stimulation electrode implant in the MFB (right). **B. Individual mice NAc DA release for rewards and omissions in Cplx.** Each line and colour is an individual mouse, averaged for all trials during last Cplx session, in [-3s:3s] time window locked on location entry. Every single mouse included in the results displayed reward-induced peaks and omission-induced dips of DA release significantly different from zero (dashed black lines). **C-D. DA cell activity using GCaMP fiber photometry.** DATiCre mice were injected with an AAV to express either GCaMP6f or GCaMP7c in VTA DA neurons, implanted with an optic fiber in the VTA, and stimulation electrode in the MFB, to assess DA neuron activity in the task. **C. GCaMP6f.** Using similar experimental procedures and signal analyses in the Cplx context, calcium dynamics of VTA DA neurons show similar reward and omission-induced transients than NAc lateral shell DA release, in this case with smaller transients amplitudes (worse signal-to-noise-ratio) whether positive or negative, and faster kinetics for positive transients. **D. GCaMP7c.** Same as B for GCaMP7c, with greater transient amplitudes (better signal-to-noise-ratio) and slower kinetic, whether positive or negative. (In B, C, D, curves are shown as mean ±sem for a single session, n is the number of reward or omission trials in this session.)

**Fig. S5:**
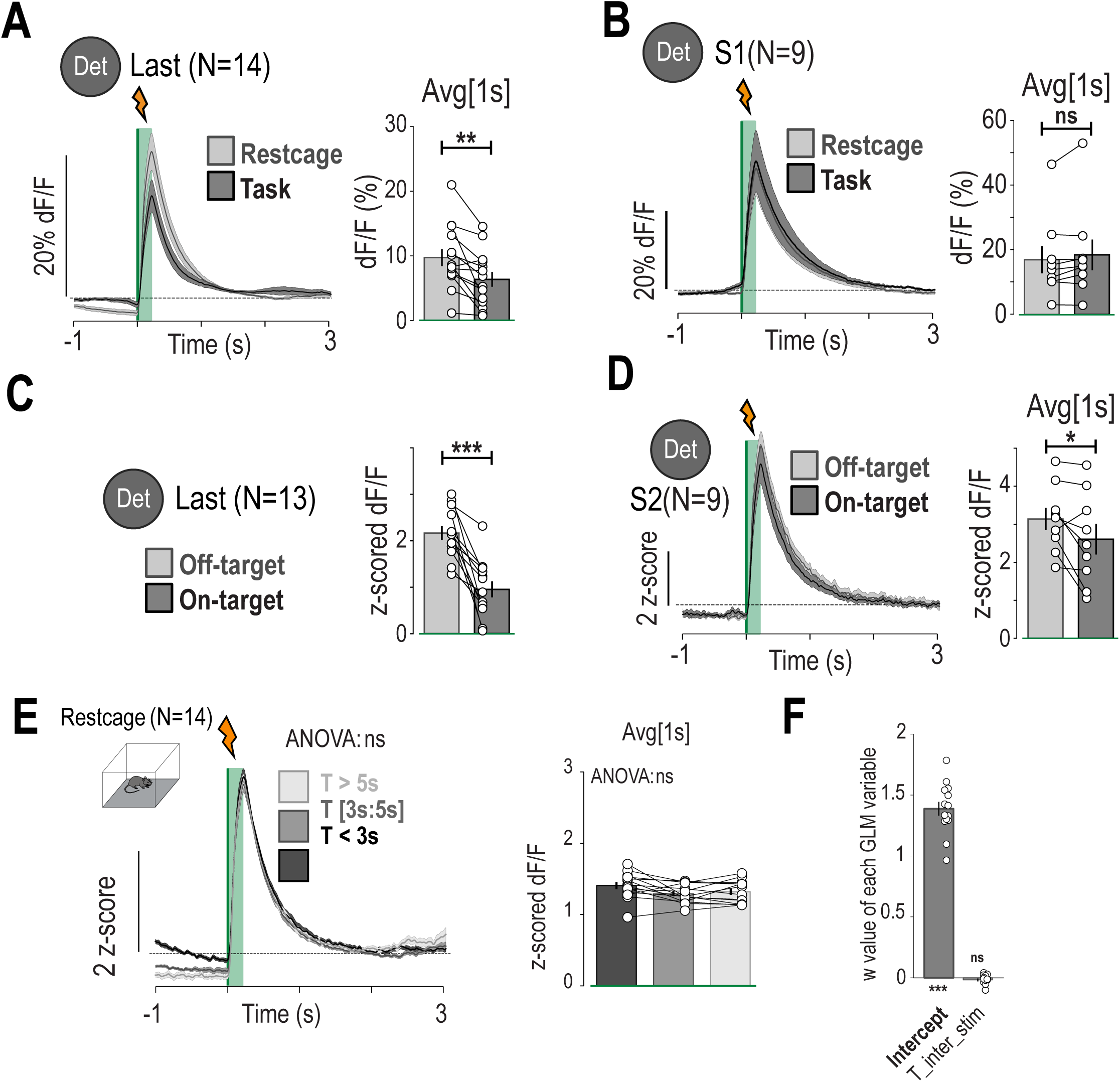
DA fiber photometry signals in various configurations. **A.** DA response to expected (Task) vs unexpected (Restcage) ICSS (same current intensity) in Det Last session. **B.** Same as A but during first session (S1) of conditioning in Det. **C.** DA response to expected (On-target) vs unexpected (Off-target) ICSS (same current intensity) in Det Last session. **D.** Same as C but during second session (S2) of conditioning in Det. **E.** Comparison of DA response for short (<3s), mid ([3s:5s]) and long (>5s) intervals of stimulation in restcage. Curves are shown as mean ±sem for session-wise average of several mice, N is the number of mice in each condition. Bar plots are shown as mean ±sem, in addition to individual data points. **F.** GLM of the amplitude of DA response as a function of interval of stimulation in the restcage, the same day as Last Det. Weight value of each variable compared to zero.

**Fig. S6:**
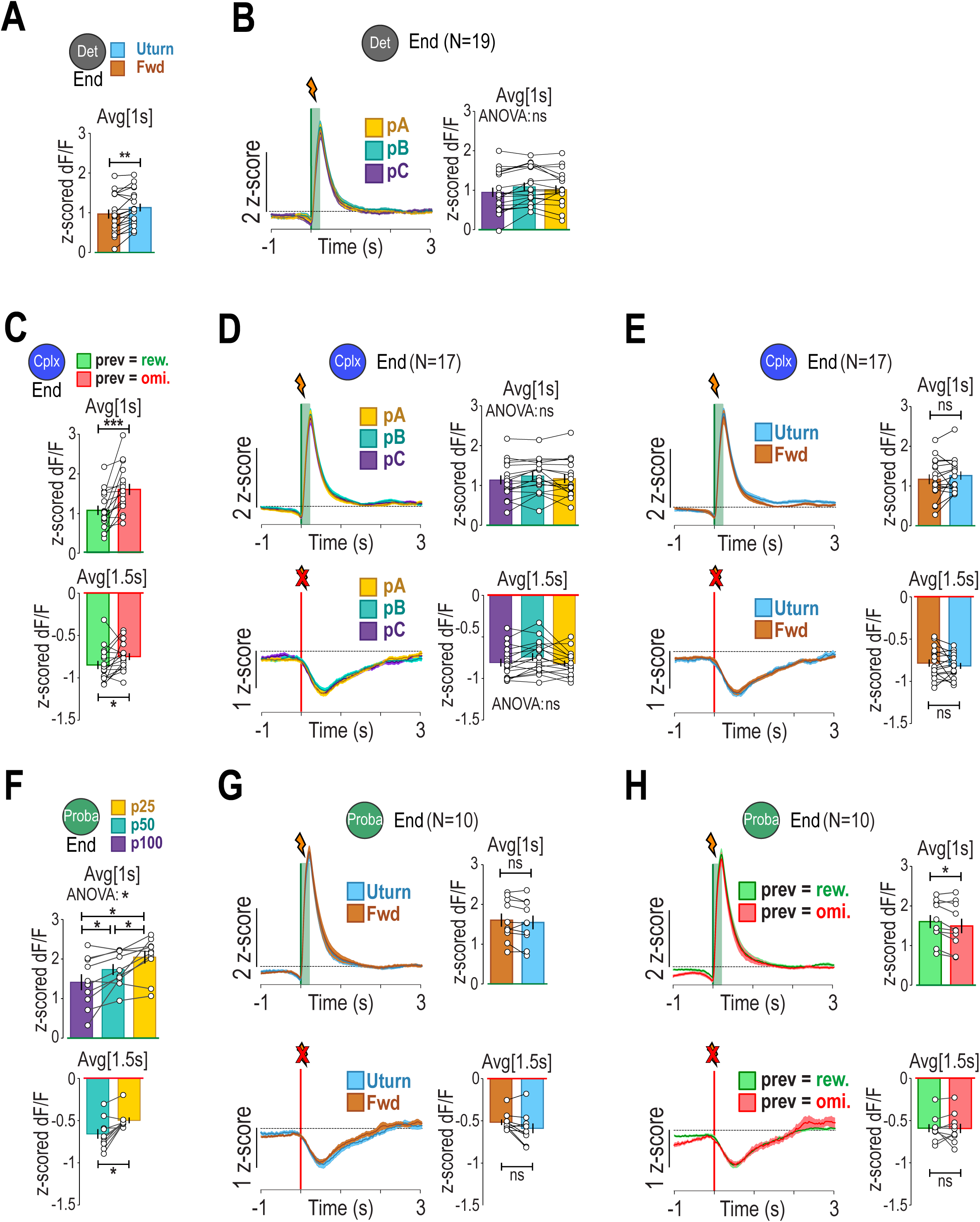
Additional analyses of NAc DA release regarding task behavioral features. **A.** Individual data corresponding to transient peak of Fig 4E. Comparison between Fwd and Uturn. **B.** Target effect on DA transients in Det End. Comparison between pA, pB and pC. **C.** Previous outcome effect on DA transients in Cplx End. Individual data corresponding to Fig 4F. **(Top)** Reward-induced transients comparison between previous reward and previous omission. **(Bottom)** Omission-induced transients comparison between previous reward and previous omission. **D.** Target effect on DA transients in Cplx End. (Top) Reward-induced transients comparison between pA, pB and pC. **(Bottom)** Omission-induced transients comparison between pA, pB and pC. **E.** Trajectory effect on DA transients in Cplx End. **(Top)** Reward-induced transients comparison between Uturn and Fwd. **(Bottom)** Omission-induced transients comparison between Uturn and Fwd. **F.** Target effect on DA transients in Proba End. Individual data corresponding to Fig 4G **(Top)** Reward-induced transients comparison between p100, p50 and p25. **(Bottom)** Omission-induced transients comparison between p50 and p25. **G.** Trajectory effect on DA transients in Proba End. **(Top)** Reward-induced transients comparison between Uturn and Fwd. **(Bottom)** Omission-induced transients comparison between Uturn and Fwd. **H.** Previous outcome effect on DA transients in Proba End. **(Top)** Reward-induced transients comparison between previous reward and previous omission. **(Bottom)** Omission-induced transients comparison between previous reward and previous omission. (Bar plots are shown as mean ±sem, in addition to individual data points. Curves are shown as mean ±sem. N is always the number of mice in each context.)

**Fig. S7:**
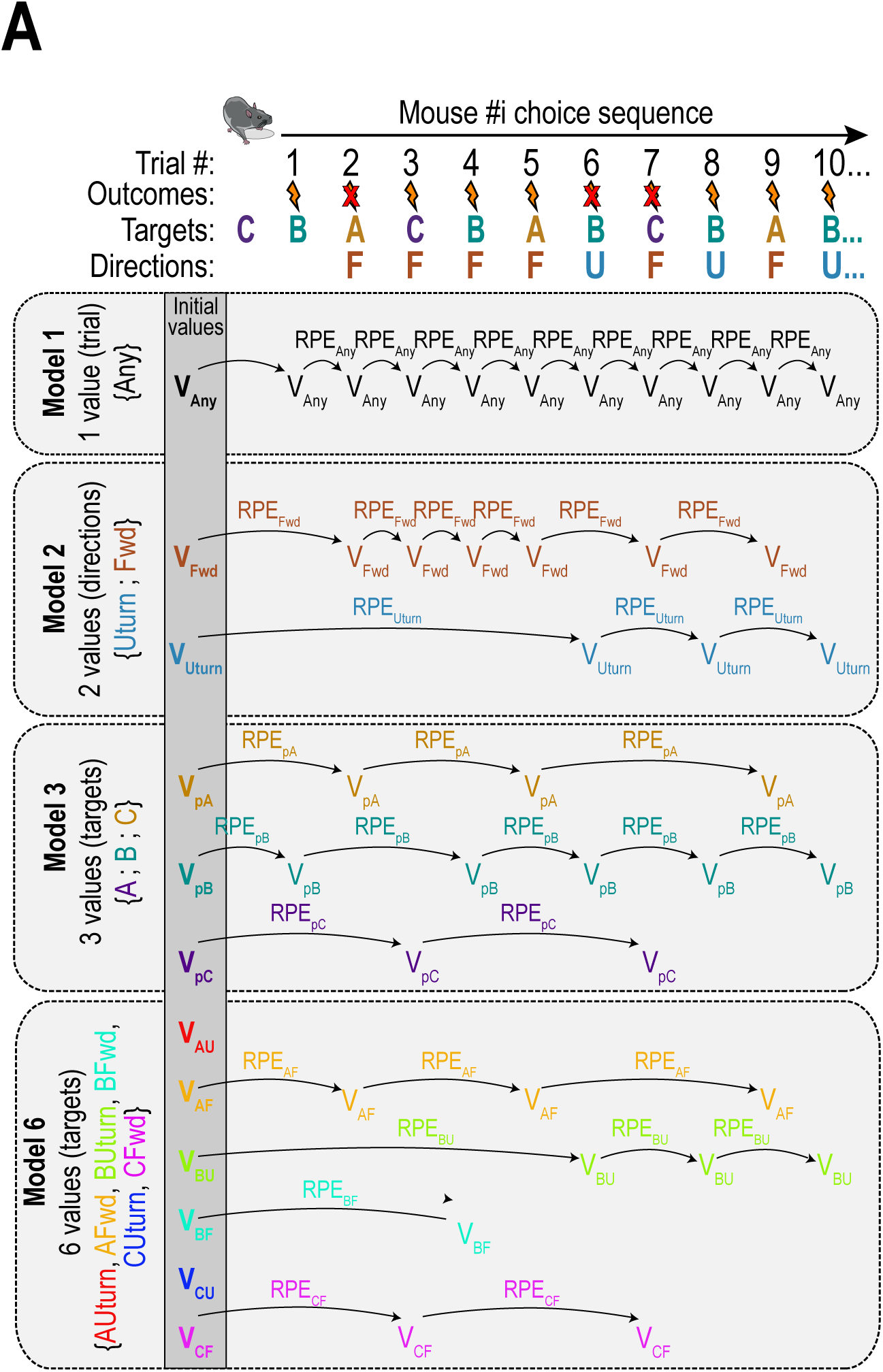
Detailed schematic of RL modelling for each of the four models. From actual mice choice sequences, we applied RL models and computed corresponding RPEs. The first model consists in single value representation “going to any target” or “performing any trial” to get a reward, where we simply compute V_expected_ = { V_Any_ } and RPE_Any_ at each trial. The second model consists in two value representations depending on chosen trajectory V_expected_ = { V_Fwd_; V_Uturn_ }. In this case, RPE_Uturn_ and RPE_Fwd_ are specific and computed separately for each of those two actions. The third model consists in three value representations depending on chosen target V_expected_ = { V_pA_; V_pB_; V_pC_ }. Again, RPE_pA_, RPE_pB_ and RPE_pC_ are computed for each target independently. The fourth model consist in six value considering forward and Uturn for each position A, B or C V_expected_ = { V_AF_; V_AU_; V_BF_; V_BU_; V_CF_; V_CU_ }. Again, RPE_AF_, RPE_AU_, RPE_BF_, RPE_BU_ RPE_CF_, and RPE_CU_ are computed for each target independently.

**Fig. S8:**
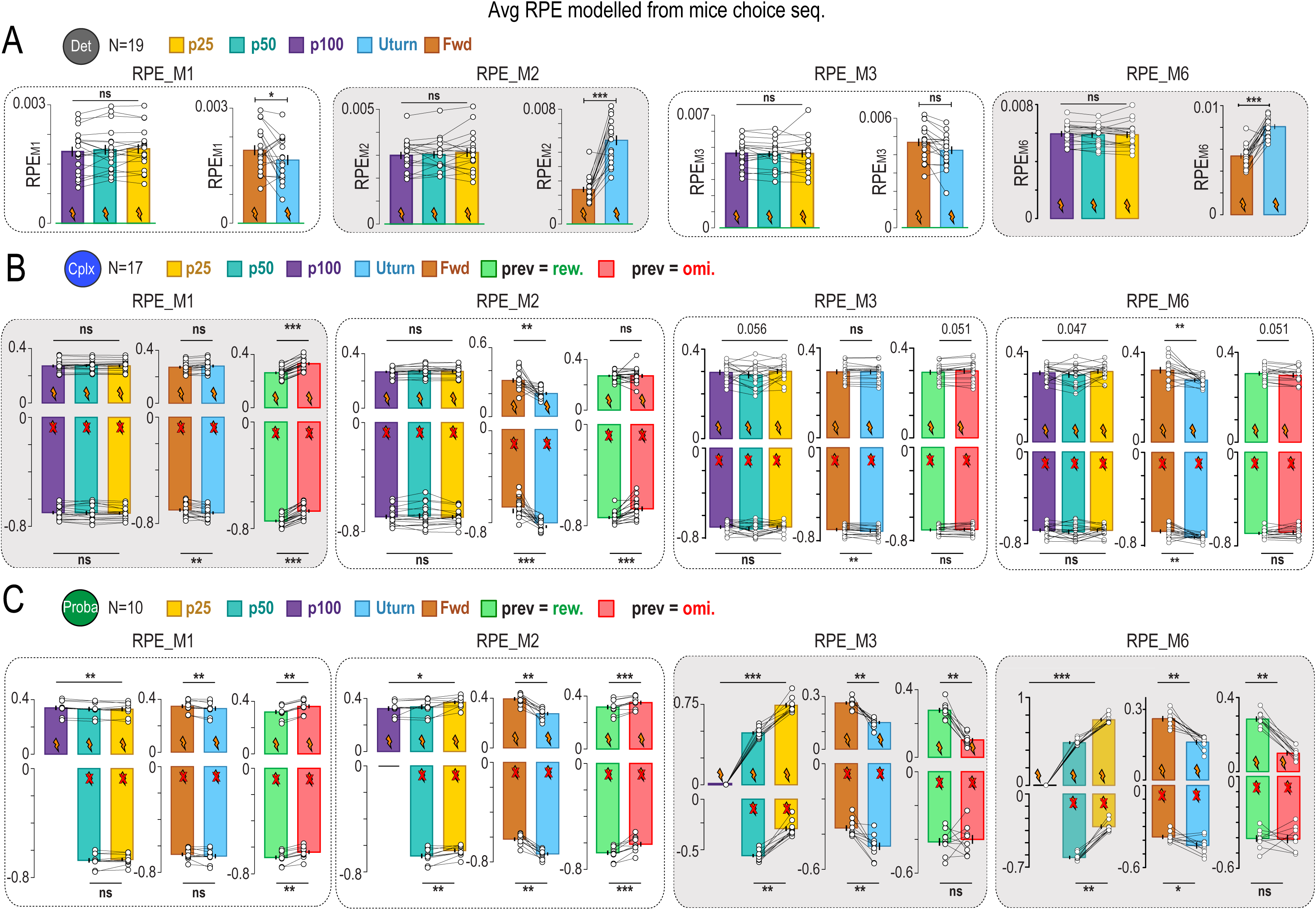
Additional information on the four RL models, comparison of computed RPEs in various behavioral scenarios, and results in Proba Change context. **A-C. For each model (M_1_ to M_6_), computed RPEs were averaged over mice sessions in the same scenarios used to characterize DA responses (regarding target, trajectory, and previous outcome).** The model that qualitatively reproduces best DA responses in all scenarios in given context is supposed to be the best value representation that mice are using in this context. **A. End Det context.** Average M1, M2, M3 and M6-computed RPE comparison between targets, and trajectories. **B. End Cplx context. From top to bottom:** Average M1, M2, M3 and M6-computed RPE comparison between targets, trajectories and previous outcome. **C. End Proba context. From top to bottom:** Average M1, M2, M3 and M6-computed RPE comparison between targets, trajectories and previous outcome. Bar plots are shown as mean ± sem, in addition to individual data points. N is always the number of mice in each context. Some of this plots are also shown in Figure 5

**Fig. S9:**
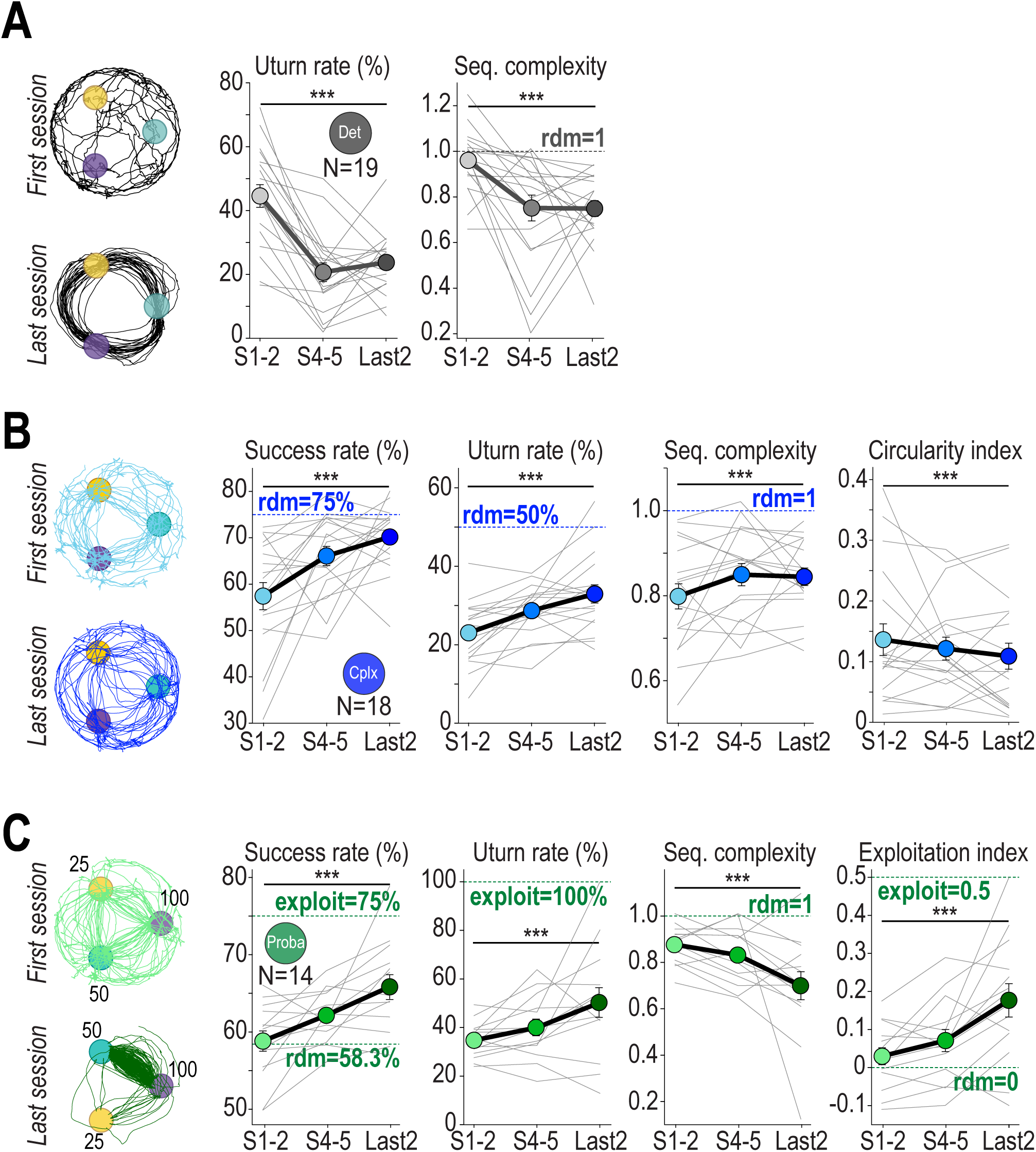
Evolution of choice parameters across rules and sessions in GRAB DA mice. **A. Evolution of choice parameters across Det sessions. Left:** Example trajectories of first and last Det session. Comparison of **(middle)** Uturn rate and **(right)** sequence complexity between sessions 1&2, sessions 4&5 and last 2 sessions in Grab-DA mice. **B. Evolution of choice parameters across Cplx sessions. Left:** Example trajectories of first (cyan) and last (blue) Cplx sessions. Comparison of **(middle-left)** Success rate, **(middle)** Uturn rate, **(middle-right)** sequence complexity and **(right)** circularity index between sessions 1&2, sessions 4&5 and last 2 sessions in Grab-DA mice. **C. Evolution of choice parameters across Proba sessions. Left:** Example trajectories of first (light green) and last (dark green) Proba sessions. Comparison of **(middle-left)** Success rate, **(middle)** Uturn rate, **(middle-right)** sequence complexity and **(right)** exploitation index between sessions 1&2, sessions 4&5 and last 2 sessions in Grab-DA mice. (data are shown as mean ±sem, in addition to individual data points. In D, each data point is one animal at one time point. signal curves are shown as mean ±sem. In A, data are shown as mean ±sem for clarity. Individual data points are available upon requests. Due to multiple corrections generating dilutions in p-values, **#** symbol has been used in the figure to highlight p<0.12 after correction. N is always the number of mice in each context.)

**Fig. S10:**
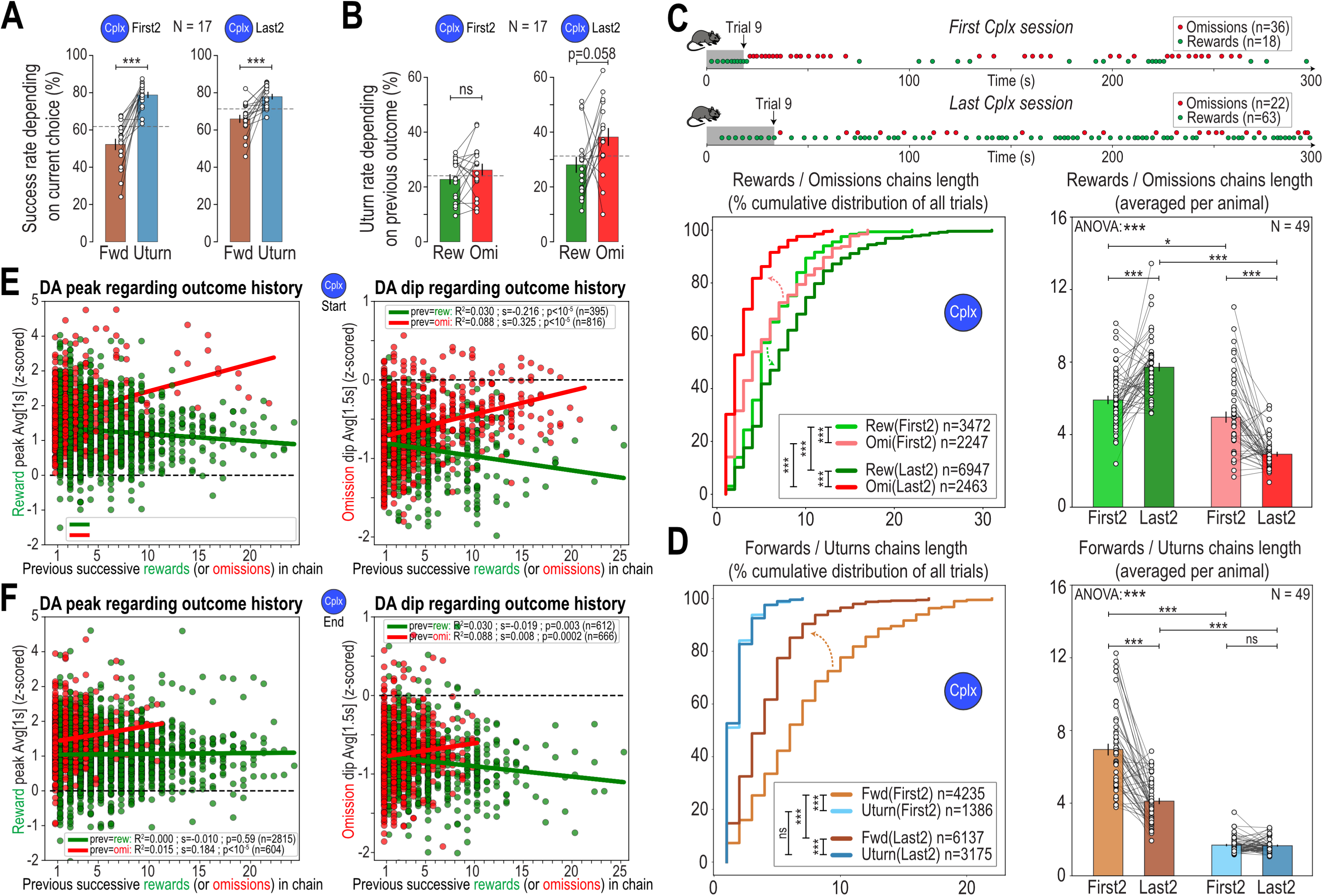
Additional analyses of DA transients and choice behavior in Cplx. **A. Success rate depending on current Uturn/Fwd choice in Cplx. Left:** First 2 sessions. **Right:** Last 2 sessions. **B. Uturn rate depending on previous outcome in Cplx. Left:** First 2 sessions. **Right:** Last 2 sessions. **C. Analysis of chains of successive rewards and omissions in Cplx. Top:** In early Cplx, mice tend to keep repeating circular patterns and therefore get long series of omissions. In late Cplx, omissions are regularly distributed, generating smaller chains, as expected from a random agent. **Bottom-left:** Cumulative distribution of reward and omission chain lengths during first 2 and last 2 Cplx sessions. **Bottom-right:** Average chain lengths per mouse. **D. Same for chains of successive forwards and Uturns. Left:** Cumulative distribution of forward and Uturn chains length during first 2 and last 2 Cplx sessions. **Right:** Average chains length per mouse. For regressions in A, B, each dot is a trial of one mouse. In C, D, E, F, Bar plots are shown as mean±sem, in addition to individual data points. In E, F, cumulative distribution are computed for all trials of all mice together. n is always the number of trials, and N the number of mice, in each context.) **E. Linear regressions of DA transients depending on the number of successive previous rewards or omissions in chains in Cplx Start. Left:** Reward-induced DA peak amplitudes regarding length of successive previous rewards chains (green) or omissions chains (red). **Right:** Same for omission-induced DA dip amplitudes regarding length of successive previous rewards chains (green) or omissions chains (red). **F. Same for Cplx End. Left:** Reward-induced DA peak amplitudes regarding length of successive previous rewards chains (green) or omissions chains (red). **Right:** Same for omission-induced DA dips amplitude regarding length of successive previous rewards chains (green) or omissions chains (red).

**Fig. S11:**
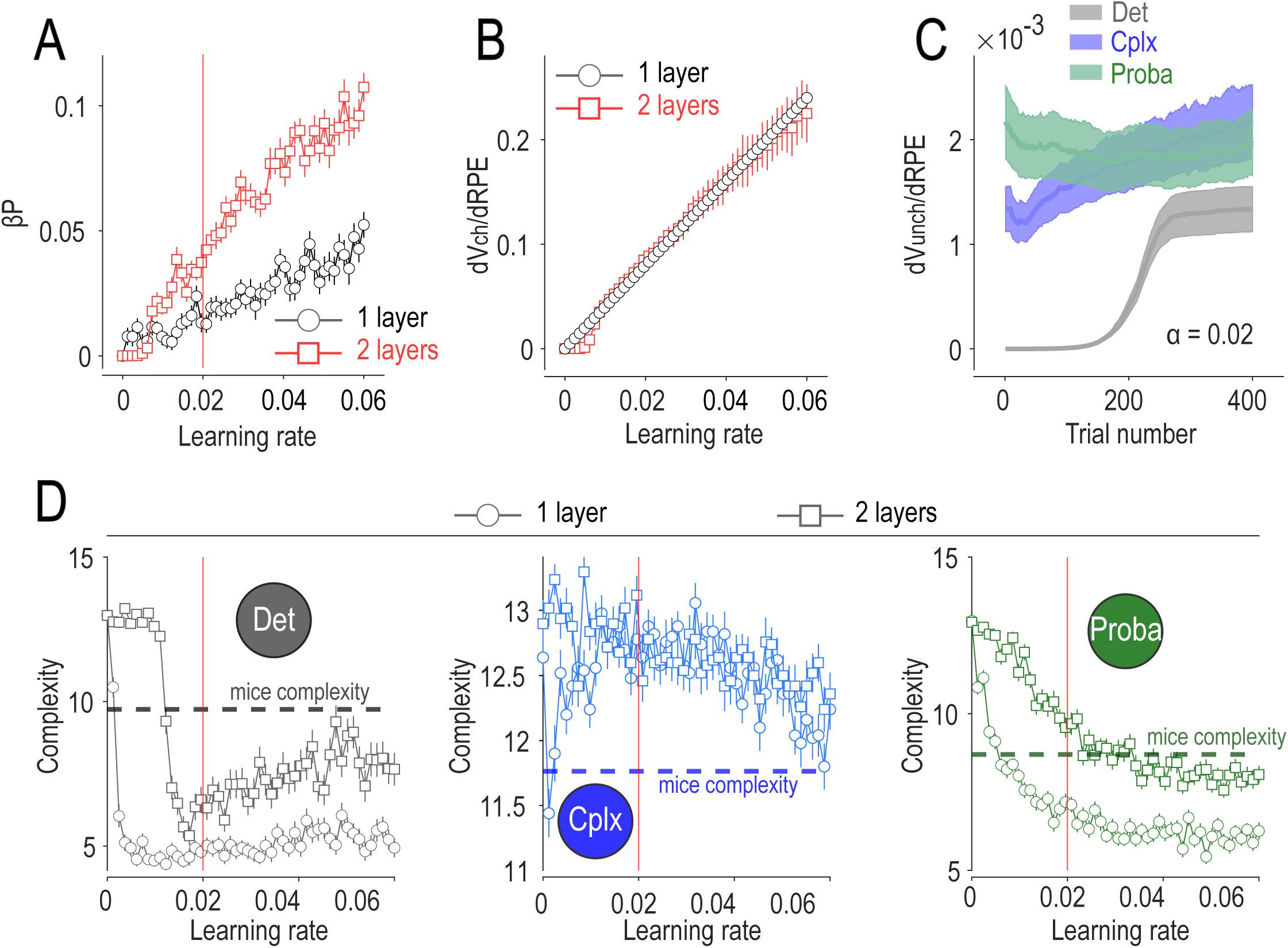
Property and performance comparison between 1-layer and 2-layers models. **A.** Sensitivity of model’s RPE to previous outcome in Cplx (βP of the full linear model, mean±SEM) according to imposed learning rate (α parameter). The vertical red bar shows the learning rate used in the rest of the paper (α = 0.02). **B.** Derivate of Value at t+1 of the action chosen at t according to RPE at t, calculated for different learning rate (mean±SEM). This figure shows that the difference in between the 2 models in A is not explained by a difference in the amplitude of the update for a given RPE between the 2 models. **C.** Evolution of the derivate of Value at t+1 of the actions unchosen at t according to RPE at t, for the learning rate used in the paper (α = 0.02), for the 2 layers model. The derivate evolves to become strictly positive through training schedule, while this derivate is 0 by definition for the 1 layer model, thus providing explanation for the difference seen in A. **D.** Mean complexity of behavior of the 2 models, calculated on a sequence of 50 trials, accross the 3 rules. This measure provides an estimation of how stereotyped (repetition of the same choices) the behavior of each model is. The dotted line shows the same estimation on the actual mice data. The 1-layer model tends to produce very stereotyped behavior (in Det and Proba) while the 2-layers model is closer to the actual data.

## Notes

### Competing Interest Statement

The authors have declared no competing interest.

### Summary of Updates

This revised version adds a neural-network reinforcement-learning (deep RL) agent to complement the interpretable RL/RPE analyses. A single deep RL agent trained by temporal-difference learning reproduces the rule-specific behavioral policies and recapitulates the rule-dependent structure of dopamine-like prediction-error signals, resulting in new/updated main figures (Figs. 6-7) and associated Methods and Supplementary analyses. We also updated the title, text and author list/affiliations accordingly.

